# Neuronal constituents and putative interactions within the *Drosophila* ellipsoid body neuropil

**DOI:** 10.1101/394833

**Authors:** Jaison Jiro Omoto, Bao-Chau Minh Nguyen, Pratyush Kandimalla, Jennifer Kelly Lovick, Jeffrey Michael Donlea, Volker Hartenstein

## Abstract

The central complex (CX) is a midline-situated collection of neuropil compartments in the arthropod central brain, implicated in higher-order processes such as goal-directed navigation. Here, we provide a systematic genetic-neuroanatomical analysis of the ellipsoid body (EB), a compartment which represents a major afferent portal of the *Drosophila* CX. The neuropil volume of the EB, along with its prominent input compartment, called the bulb, is subdivided into precisely tessellated domains, distinguishable based on intensity of the global marker DN-cadherin. EB tangential elements (so-called ring neurons), most of which are derived from the DALv2 neuroblast lineage, interconnect the bulb and EB domains in a topographically-organized fashion. Using the DN-cadherin domains as a framework, we first characterized the bulb-EB connectivity by Gal4 driver lines expressed in different DALv2 ring neuron (R-neuron) subclasses. We identified 11 subclasses, 6 of which correspond to previously described projection patterns, and 5 novel patterns. These subclasses both spatially (based on EB innervation pattern) and numerically (cell counts) summate to the total EB volume and R-neuron cell number, suggesting that our compilation of R-neuron subclasses approaches completion. EB columnar elements, as well as non-DALv2 derived extrinsic ring neurons (ExR-neurons), were also incorporated into this anatomical framework. Finally, we addressed the connectivity between R-neurons and their targets, using the anterograde trans-synaptic labeling method, *trans*-Tango. This study demonstrates putative interactions of R-neuron subclasses and reveals general principles of information flow within the EB network. Our work will facilitate the generation and testing of hypotheses regarding circuit interactions within the EB and the rest of the CX.

## Introduction

The central complex (CX) is an evolutionarily conserved, higher-order neuropil in the arthropod brain thought to integrate sensory and motor information to coordinate and maintain locomotor behavior, thus enabling appropriate navigation. *Drosophila* mutations that produce structural abnormalities in CX neuropils result in flies with deficiencies in walking and flight (Martin et al., 1999; Strauss and Heisenberg, 1993). More targeted manipulations, such as silencing of specific CX neuron subclasses, compromise vision-based memories associated with spatial orientation and location (Neuser et al., 2008; Ofstad et al., 2011). Similar themes emerge from anatomical, electrophysiological, and behavioral studies investigating the CX in other insects. In the cockroach CX, for example, single unit activity correlated with changes in locomotor intensity, turning behavior, or heading direction have been identified (Bender et al., 2010; Guo and Ritzmann, 2013; Varga and Ritzmann, 2016). In addition, electrical stimulation of CX neurons in the freely walking cockroach has yielded direct evidence linking CX activity to downstream locomotor output (Martin et al., 2015). In other insects, such as locust, cricket, monarch butterfly, and dung beetle, neurons in the CX are tuned to celestial visual cues such as the sun or pattern of polarized skylight. These cues provide the stable environmental signals required to accurately derive relative heading information for short or long range navigations (Heinze and Homberg, 2007; Heinze and Reppert, 2011; Jundi et al., 2014; 2015).

The CX consists of four neuropil compartments: the upper (CBU) and lower (CBL) halves of the central body (CB), protocerebral bridge (PB), and paired noduli (NO) (Hanesch et al., 1989; Ito et al., 2014; Strausfeld, 2012). In *Drosophila*, the upper and lower halves of the central body are designated as the fan-shaped body (FB) and ellipsoid body (EB), respectively (Fig.1A). These neuropil compartments are formed by two orthogonally arranged neuronal populations: (1) columnar (small-field) neurons which interconnect the CX compartments along the antero-posterior axis; (2) tangential (large-field) neurons which provide input from lateral brain neuropils to the CX (Fig.1B, C). Terminal arborizations of these neurons define distinct vertical columns and horizontal layers that can be visualized by markers for synaptic or cell adhesion proteins that globally label, but exhibit variable density in, the neuropil. Based on Bruchpilot immunostaining, seven layers were identified in the *Drosophila* CBU (=FB; Fig.1A) (Wolff et al., 2015). The CBL (=EB) also exhibits a layered organization (Pfeiffer and Homberg, 2014). In *Drosophila*, this compartment undergoes a morphogenetic transformation during pupal development, whereby the lateral ends of the originally bar-shaped EB primordium bend ventrally to adopt a toroidal arrangement (Lovick et al., 2017; Xie et al., 2017; Young and Armstrong, 2010a; Fig.1A). As a result, tangential neurons of the EB display a circular shape, and hence were called “ring neurons” (Hanesch et al., 1989; Fig.1C). Likewise, layers within the EB are annuli, rather than horizontal slabs (Fig.1A). Based on labeling with DN-cadherin, we have defined five distinct annular domains, termed anterior (EBa), inner and outer central (EBic and EBoc), and inner and outer posterior (EBip and EBop) domains (Omoto et al., 2017; Fig.1D-G).

**Figure 1.**
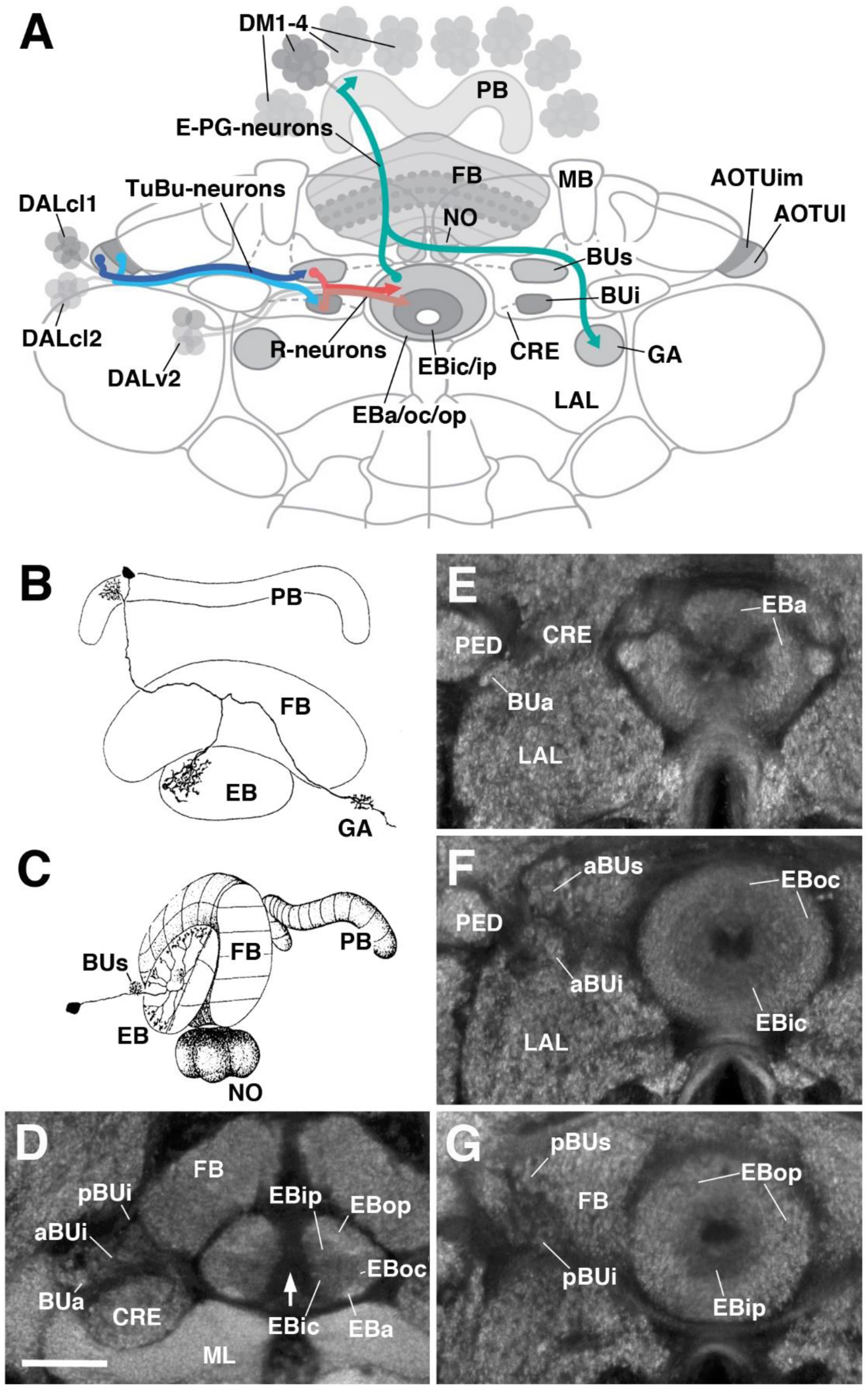
General overview of the ellipsoid body: neuronal interactions and compartmentalization (A) Schematized overview of interacting neuronal populations of the ellipsoid body. Gray indicates relevant lineages, neuron types, and neuropil compartments. The anterior visual pathway, divided into the superior and inferior bulb pathways, provides input to R-neurons. Superior bulb pathway: Tuberculo-bulbar (TuBu) neurons of lineage DALcl1 (dark blue) project from the lateral domain of the anterior optic tubercle (AOTUl) to the superior bulb (BUs), which then innervate R-neurons (dark red) that project to the anterior and outer central domains of the ellipsoid body (EBa/oc). Inferior bulb pathway: TuBu neurons of lineage DALcl2 (light blue) project from the intermediate medial domain of the anterior optic tubercle (AOTUim) to the inferior bulb (BUi), which then innervate R-neurons (light red) that project to the inner central and inner posterior domains of the ellipsoid body (EBic/ip). Columnar elements of the DM lineages (turquois), such as E-PG neurons, form recurrent circuitry interconnecting the protocerebral bridge (PB), EB, and gall (GA) of the lateral accessory lobe (LAL). Many other neuron types not shown interconnect the fan-shaped body (FB) and noduli (NO) as well. (B and C) Representative examples of columnar and tangential elements of the ellipsoid body. (B) E-PG neuron and (C) R2-neuron. Images adapted from Hanesch et al., 1989. (D-G) Confocal z-projections illustrating domains of the bulb and ellipsoid body, visible with DN-cadherin (DNcad) immunostaining (gray). (D) horizontal section; (E-G) frontal sections at three different antero-posterior depths. (D) Horizontal section (anterior pointing downward) depicting the length of the ellipsoid body canal (arrow). The EB is situated within an indentation of the FB, located posteriorly. All five EB domains, distinguishable based on DNcad expression levels, are visible: High intensity staining in the anterior-most part of the EB defines the anterior domain (EBa). Posterior to EBa is the inner central domain (EBic) with lower DNcad signal, located medially adjacent to the outer central domain (EBoc) with moderate DNcad signal. Furthest posterior are the inner posterior (EBip) and outer posterior (EBop) domains with low and high intensity DNcad signal, respectively. In this horizontal section, the anterior bulb (BUa) as well as the anterior and posterior regions of the inferior bulb (a/pBUi) are visible, but the superior bulb, located more dorsally, is not. (E) Anterior section: the anterior bulb (BUa) and anterior domain of the EB (EBa) are visible. In addition, the anterior-most part of inner central domain (EBic) is also visible (low intensity region proximal to the EB canal). (F) Intermediate section: the anterior regions of the superior bulb (aBUs) and inferior bulb (aBUi), as well as the inner central (EBic) and outer central (EBoc) domains of the ellipsoid body are visible. (G) Posterior section: the posterior regions of the superior bulb (pBUs) and inferior bulb (pBUi), as well as the inner posterior (EBip) and outer posterior (EBop) domains of the ellipsoid body are visible. Other abbreviations: CRE, crepine; MB, mushroom body; ML and PED, medial lobe and peduncle of the mushroom body, respectively.

Clonal studies in *Drosophila* show that the neuronal architecture of the CX is organized into lineage-based modules (Ito and Awasaki, 2008; Yang et al., 2013), a ground plan that is likely conserved across insects (Boyan et al., 2017). A lineage refers to the set of sibling neurons derived from an individual neural progenitor called a neuroblast, and the entire central brain is generated from a fixed number of approximately 100 of such neuroblasts. Four lineages (DM1-4; Fig.1A) give rise to the large number of columnar neurons of the CX (Ito and Awasaki, 2008; Yang et al., 2013). The great diversity observed among these neurons is achieved via temporal patterning of molecular determinants in dividing progenitors (Bayraktar and Doe, 2013; Doe, 2017; Wang et al., 2014). Lineages giving rise to the tangential neurons of the CX have been characterized morphologically (Ito et al., 2013; Larsen et al., 2009; Spindler and Hartenstein, 2010; Wong et al., 2013; Yang et al., 2013; Yu et al., 2013), but have not yet received much attention experimentally. The most notable exception is lineage DALv2/EBa1 (henceforth called DALv2), that generates ring neurons of the EB (Neuser et al., 2008; Omoto et al., 2017; Seelig and Jayaraman, 2013; Fig.1A). Ring neurons project their axons to distinct annular domains of the EB, and typically possess short globular dendrites (“microglomeruli”) in the bulb (BU), a neuropil compartment located laterally adjacent to the EB (Fig.1A). The bulb encompasses three main partitions (anterior, superior, inferior) that are associated with different annular domains of the EB (Fig.1A, E-G). Input to the bulb is provided by neurons of two additional lineages, DALcl1 and DALcl2 (also called AOTUv3 and AOTUv4, respectively) (Omoto et al., 2017; Wong et al., 2013; Yang et al., 2013; Yu et al., 2013). As part of the anterior visual pathway, DALcl1/2 form so-called tubercular-bulbar (TuBu) neurons which project from the anterior optic tubercle to the bulb, relaying visual information to ring neurons and thereby the CX as a whole. TuBu neurons form two lineally-segregated parallel channels, with DALcl1 establishing connections with ring neurons located in the peripheral domain of the EB via the superior bulb, and DALcl2 with central ring neurons via the inferior bulb (Omoto et al., 2017; Shiozaki and Kazama, 2017; Fig.1A).

Detailed functional studies are beginning to shed light on the circuitry involving ring neurons and their TuBu afferents and columnar efferents. Two-photon calcium imaging has revealed a discrete focus of neural activity, or “bump”, within a population of columnar neurons (“E-PGs”) that interconnect the EB, PB, and gall of the LAL. E-PG neurons encode an internal compass representation via the activity bump, which dynamically tracks the fly’s heading (Seelig and Jayaraman, 2015; Fig.1A). Additional columnar neuron populations that interconnect the PB, EB, and NO, called P-EN neurons, compute the animals’ heading by controlling the movement of the bump in the clockwise or counter-clockwise direction (Green et al., 2017; Turner-Evans et al., 2017). These findings suggest that the EB may operate as a critical hub in the CX, acting as an interface between neurons that transmit and distribute sensory information (TuBu and ring neurons), and circuits that encode and update a representation of heading direction (E-PG and P-EN neurons). In addition, internal state information is likely integrated into the EB network by additional ring neurons subclasses that signal physiological needs such as sleep and hunger drive (Dus et al., 2013; Liu et al., 2016; Park et al., 2016).

To make further inroads in understanding how the EB circuitry operates, a comprehensive knowledge of ring neurons and their upstream and downstream connectivity is required. Ultimately, a comprehensive analysis of single cells and their synaptic contacts on the light and electron microscopy level will yield complete coverage of the EB wiring diagram, and certainly inform our understanding of how EB-related computations are implemented (Zheng et al., 2018). However, a current description of subclass-specific projection patterns using genetic driver lines provides a framework to posit inter-class neural interactions that can then be tested physiologically and/or behaviorally, and will assist future efforts for such high-resolution anatomical maps. To this end, we sought to expand on previous works using this genetic-anatomical approach to more thoroughly describe the EB neuropil (Martín-Peña et al., 2014; Omoto et al., 2017; Renn et al., 1999; Young and Armstrong, 2010b). Gal4 driver lines that label ring neuron subclasses were screened and subsequently distinguished from each other based on defined criteria. Many drivers label populations corresponding to previously identified ring neuron subclasses, in addition to several, yet uncharacterized populations. The novel subclasses were given new names per the historical nomenclature system. Columnar elements were also incorporated into this anatomical framework. Based on the domain innervation pattern of each line, putative interactions between elements within the EB network are proposed. Finally, ring neuron drivers were subjected to the anterograde trans-synaptic labeling method, *trans*-Tango (Talay et al., 2017). Ring neurons occupying central domains of the EB commonly display homotypic interactions, such that neurons of a given subclass predominantly form synaptic interactions with other neurons in the same subclass. On the other hand, ring neurons occupying the peripheral domains typically display a larger degree of output into the columnar network. This highlights a fundamental difference in the connectivity, and potentially the functions, of ring neurons in different domains.

## Material and methods

### Fly lines

The following *Drosophila* Gal4 driver lines are from the Janelia Research Campus stock collection (Jenett et al., 2012), and acquired from the Bloomington *Drosophila* Stock Center (BDSC), Bloomington, Indiana: R31A12, R78B06, R80C07, R28E01, R28D01, R12G08, R35D04, R84H09, R15B07, R12B01, R59B10, R38H02, R78A01, R14G09. VT063949, VT057232, and VT011965 are Vienna Tile Gal4 driver lines (Tirian and Dickson, 2017) and were acquired from Dr. Barry Dickson. Ring neuron lines were typically identified by visually screening the Janelia FlyLight database (http://flweb.janelia.org/cgi-bin/flew.cgi). R14G09 was identified by first visually screening the Flycircuit database (http://www.flycircuit.tw/) for EB innervating neurons, yielding clone ID# VGlut-F-300355 (Chiang et al., 2011), followed by use of the NBLAST (Flycircuit to Gal4 query) search algorithm (Costa et al., 2016). Additional stocks, with citation and availability listed in parentheses: 189Y, c42, c232, c105, c507 (Renn et al., 1999; BDSC), EB1-Gal4 (Young and Armstrong, 2010b), Poxn-Gal4 (Boll and Noll, 2002; provided by Dr. H. Reichert), 10xUAS-mCD8::GFP (BDSC), TPH-Gal4 (Park et al., 2006; provided by Dr. M. Frye), TH-Gal4 (Friggi-Grelin et al., 2003; BDSC), UAS-DenMark::mCherry, UAS-syt.EGFP (Nicolaï et al., 2010; BDSC), su(Hw)attP8:HA_V5_FLAG_1 (Nern et al., 2015; BDSC), *trans*-Tango (Talay et al., 2017; provided by Dr. G Barnea).

### Clonal analysis

Mosaic analysis with a repressible cell marker (MARCM) was conducted to generate heat shock inducible, single-cell clones of ring neurons (Lee and Luo, 1999). Flies of the following genotypes were utilized: hsflp/+; FRTG13, UAS-mCD8GFP/FRTG13, tub-GAL80; tub-Gal4/+ or FRT19A, tub-GAL80, hsflp, UAS-mCD8GFP/elav^C155^-Gal4, FRT19A; UAS-mCD8GFP/+. GFP-labeled adult single cell MARCM clones were induced at the late first instar/early second instar stage by heat-shocking in a water bath at 38 °C for 30-60 min. Larvae were collected after hatching, reared at 18 °C, and heat-shocked at different time intervals between 12-144 hours (double time; corresponding to roughly 6-72h at 25 °C). Heat-shocked larvae were grown to adulthood for subsequent dissection and analysis. Single-cell analysis of TH-Gal4 positive neurons was conducted using the multicolor flip-out method (MCFO) described previously (Nern et al., 2015; Wolff et al., 2015).

### Immunostaining

3-8 day old female adults were used for all experiments; potential sexual dimorphism would be undetected in this study. Flies were grown at 25ºC on standard fly media, in low density bottles on a 12hr:12hr light/dark schedule. Immunohistochemical procedures were conducted as follows, and are similar to those previously described (Omoto et al., 2017). Adult brains were dissected in phosphate buffered saline (PBS), pH 7.4. Brains were (1) fixed in ice-cold PBS containing 4% EM-grade paraformaldehyde for 2.5-3 hours; (2) washed 4X for 15 min each with ice-cold PBS; (3) subjected to cold ethanol-PBS dehydration (five minute washes in 5, 10, 20, 50, 70, 100% EtOH); (4) stored in ‒20^°^C overnight; (5) rehydrated using the same cold EtOH series in reverse order; (6) washed 2X 15 min in cold PBS and 2X 15 min washes in cold 0.3% PBT (PBS containing 0.3% Triton X-100); (7) washed in room temperature (RT) 0.3% PBT 4X 15 min; (8) incubated in blocking buffer (10% normal goat serum in 0.3% PBT) for 30 min at RT; (9) incubated in primary antibody, diluted in blocking buffer, at 4^°^C for 3 nights; (10) washed 4X 15 min in RT 0.3% PBT; (11) incubated with secondary antibody diluted in blocking buffer at 4^°^C for an additional 3 nights; (12) washed 4X 15 min in RT 0.3% PBT and mounted using Vectashield (Vector Laboratories). For 10xUAS-mCD8::GFP panels, native fluorescence of the reporter was used to visualize the neurons. To maximally detect both pre- and post-synaptic neurons in *trans*-Tango experiments, both anti-GFP and anti-DsRed were utilized.

The following antibodies were provided by the Developmental Studies Hybridoma Bank (Iowa City, IA): rat anti-DN-cadherin (DN-EX #8, 1:20), mouse anti-Neuroglian (BP104, 1:30). Chicken anti-GFP (Abcam #ab13970, 1:1000) and Rabbit anti-DsRed (Clontech #632496, 1:1000) were also used. We also used rabbit anti-HA (1:300, Cell Signaling Technologies), and mouse anti-V5 (1:1000, Thermo Fisher Scientific).

Secondary antibodies, IgG_1_ (Jackson ImmunoResearch; Molecular Probes) were used at the following dilutions: Cy5-conjugated anti-mouse (1:300), Cy3-conjugated anti-rat (1:300). Alexa We used 488-conjugated anti-chicken (1:1000), Alexa 546-conjugated anti-rabbit (1:1000), Alexa488-conjugated anti-mouse (1:1000) from Thermo Fisher Scientific. Cy5-conjugated anti-rat (1:300) and cy3-conjugated anti-rabbit (1:300) from Abcam were also used.

### Confocal microscopy and image analysis

Samples were mounted primarily in the antero-posterior (A-P) or dorso-ventral (D-V) orientation, and in some cases the postero-anterior (P-A) orientation. D-V orientation required constructing a crevice using two closely neighboring pieces of tape followed by two cover slips, into which the brain can be inserted dorsal-side up. Whole-mounted brains were imaged using confocal microscopy [LSM 700 Imager M2 using Zen 2009 (Carl Zeiss Inc.)]. Series of optical sections were imaged using a 40X oil lens with a numerical aperture of 1.3, a zoom factor of 1.0, at 1.2-μM intervals, and 1024 x 1024 pixel resolution. Digitized images of confocal sections were processed in FIJI (Schindelin et al., 2012; http://fiji.sc/). The ellipsoid body, relative to the rest of the brain, exhibits a tilt on its frontal axis such that the ventral half is oriented anteriorly. We established standard, reproducible views of the EB in both the frontal and horizontal planes (used in all figures) by digitally tilting the z-stack using the “Interactive Stack Rotation” plugin (http://imagej.net/Interactive_Stack_Rotation). Antero-posteriorly and dorso-ventrally mounted preparations were digitally tilted such that the canal of the ellipsoid body was oriented parallel and perpendicular to the z-axis, respectively. In several cases, particularly in cases for z-projections that span large depths (greater than ~75 μM; ex. dorsal *trans*-Tango images), background labeling was manually removed in FIJI to improve visualization of entire neuronal ensembles. Cell counts were conducted manually using the FIJI “Cell Counter” plugin (https://imagej.nih.gov/ij/plugins/cell-counter.html). Cell body clusters on both sides of each brain were counted for at least three samples per driver line. Mean and standard error of the mean was calculated. Schematics were generated in Adobe Illustrator and figures constructed in Adobe Photoshop. Videos were compiled using Camtasia 9.1 with annotations on individual slices made using FIJI “Dotted Line” plugin (https://imagej.nih.gov/ij/plugins/dotted-line.html).

## Results

### Classification of EB ring neurons: criteria and general considerations

Using Golgi staining to characterize individual CX neuron types in *Drosophila*, the term “ring neuron” was coined by Hanesch et al., (1989), defined as “large-field neurons forming ring-like arborizations around the ellipsoid body canal”. Although “R-neuron” is commonly used as a synonymous abbreviation for “ring neuron”, the latter (full) term was originally used by Hanesch et al., (1989) as an umbrella designation for two major neuron types, R and ExR (“extrinsic ring neurons”). R-neurons represent the most abundant type, with cell bodies located in the anterior cell body rind (also called cortex herein), dorso-laterally of the antennal lobes. ExR-neurons were defined as ring neurons that have “extensive arborizations outside of the EB”. Due to the utility of this distinction to refer to ring neurons derived from distinct neuroblast lineages (DALv2 = R-neurons; DM4-6 and BAmv1 = ExR-neurons), we re-adopt it for this study (see below).

For the time being, we adopt and expand upon the historical *Drosophila* ring neuron nomenclature system (i.e. R1, R2, ExR1, etc.), initially introduced by Hanesch et al., (1989) with other studies largely following suit (Omoto et al., 2017; Renn et al., 1999; Young and Armstrong, 2010b). Wolff et al., (2015) developed a formal nomenclature system for neuron types of the PB, in which each cell type was named based on a unique, descriptive collection of identifiers. As more information becomes available, adopting a unified nomenclature system in conjunction with subordinate colloquial terminology, for ring neurons and other neurons comprising the rest of the CX or brain in general, may be most suitable. We propose that this prospective system would ideally incorporate lineage classification as one of these identifiers, as the fly brain is inherently organized into structurally and developmentally-defined clonal units.

In most cases, the Gal4 drivers that label the ring neurons described in this study were visually screened from the Janelia (Jenett et al., 2012) or Vienna Tiles (Tirian and Dickson, 2017) collections and subsequently stained with the global neuropil marker DN-cadherin. Drivers were classified as labeling a unique ring neuron subclass based on the following criteria: 1) the EB DN-cadherin subdomain occupied by the circular, predominantly axonal, arbors, 2) the trajectory and morphology of said projections, and 3) the location of their presumed dendritic proximal neurites, typically microglomeruli in the bulb or fibrous neurites in the lateral accessory lobe (LAL).

Altogether, we identified fifteen unique ring neuron subclasses: eleven R-neuron and four ExR-neuron subclasses. This expands the catalog from six R-neuron (R1, R2, R3, R4m, R4d, R5) and two ExR-neuron (ExR1 and ExR2) subclasses, from previous reports (Hanesch et al., 1989; Omoto et al., 2017; Renn et al., 1999; Young and Armstrong, 2010b). However, it is critical to note a caveat of this study: each driver labels a population of neurons which were not anatomically evaluated on a single-cell basis, as has been done for the neurons innervating the PB (Wolff et al., 2015). Indeed, multicolor-flip out analysis (MCFO) of some ring neuron drivers from this study yielded qualitatively distinct anatomical subtypes, even within a superficially homogenous population (data not shown). Therefore, although this study significantly expands upon the cohort of known ring neuron subclasses, in the absence of higher-resolution methods (single-cell light microscopy, TEM reconstructions) and supplementary genetic/physiological evidence, the precise diversity of ring neuron subclasses is still underestimated. Nonetheless, this study provides a more complete catalog and explicit criteria with which ring neuron subclasses can be anatomically defined. These criteria may be used as a framework to define new subclasses identified in subsequent studies.

One supplementary objective of this study is to resolve discrepancies in the literature regarding ring neuron subclasses, a consequence of somewhat undefined criteria and lack of spatial resolution. We have reevaluated previously published driver lines based on the proposed criteria and have found that oftentimes, a given ring neuron subclass has been called distinct names in different studies. Alternatively, a previously unidentified subclass has been assumed to be one of the preexisting subclasses because it appeared similar. When examining each subclass below, we will refer to pertinent examples of this, and provide data to reevaluate these comparisons based on our criteria.

### R-neurons: lineage DALv2/EBa1

The most abundant ring neuron type is the R-neurons, whose cell bodies are located dorso-laterally of the antennal lobes and exhibit projections that extend dorso-posteriorly, branching off localized neurites into the bulb (BU) or LAL, enter the lateral ellipsoid fascicle [(LE; Lovick et al., 2013; Pereanu et al., 2010; Strausfeld, 2012); also called the “isthmus tract” (Ito et al., 2014)], and via a circular process, terminates medially into the EB (Figs.2 and 3). In the EB, distal neurites of R-neurons either project centrifugally (“inside-out”; Fig.2) or centripetally (“outside-in”; Fig.3). Clonal analysis of fly brain lineages revealed a single paired type I neuroblast that generates R-neurons called DALv2 (Omoto et al., 2017; Wong et al., 2013), also called EBa1 (Ito et al., 2013; Yu et al., 2013). The driver Poxn-Gal4 (Boll and Noll, 2002; data not shown), which labels the majority of (but possibly not all) DALv2 R-neurons, can, at first approximation, be used to estimate the total number of R-neurons. Quantification of these cells reveals 158±9 R-neurons per brain hemisphere (PBH). Despite the aforementioned caveats, the following catalog of R-neuron drivers is likely close to comprehensive, considering that summation of the neurons from each R-neuron driver (11 lines) totals ~176 cells PBH (see below). In the following sections, we summarize the neuroanatomy of Gal4 lines that label unique R-neuron patterns.

**Figure 2.**
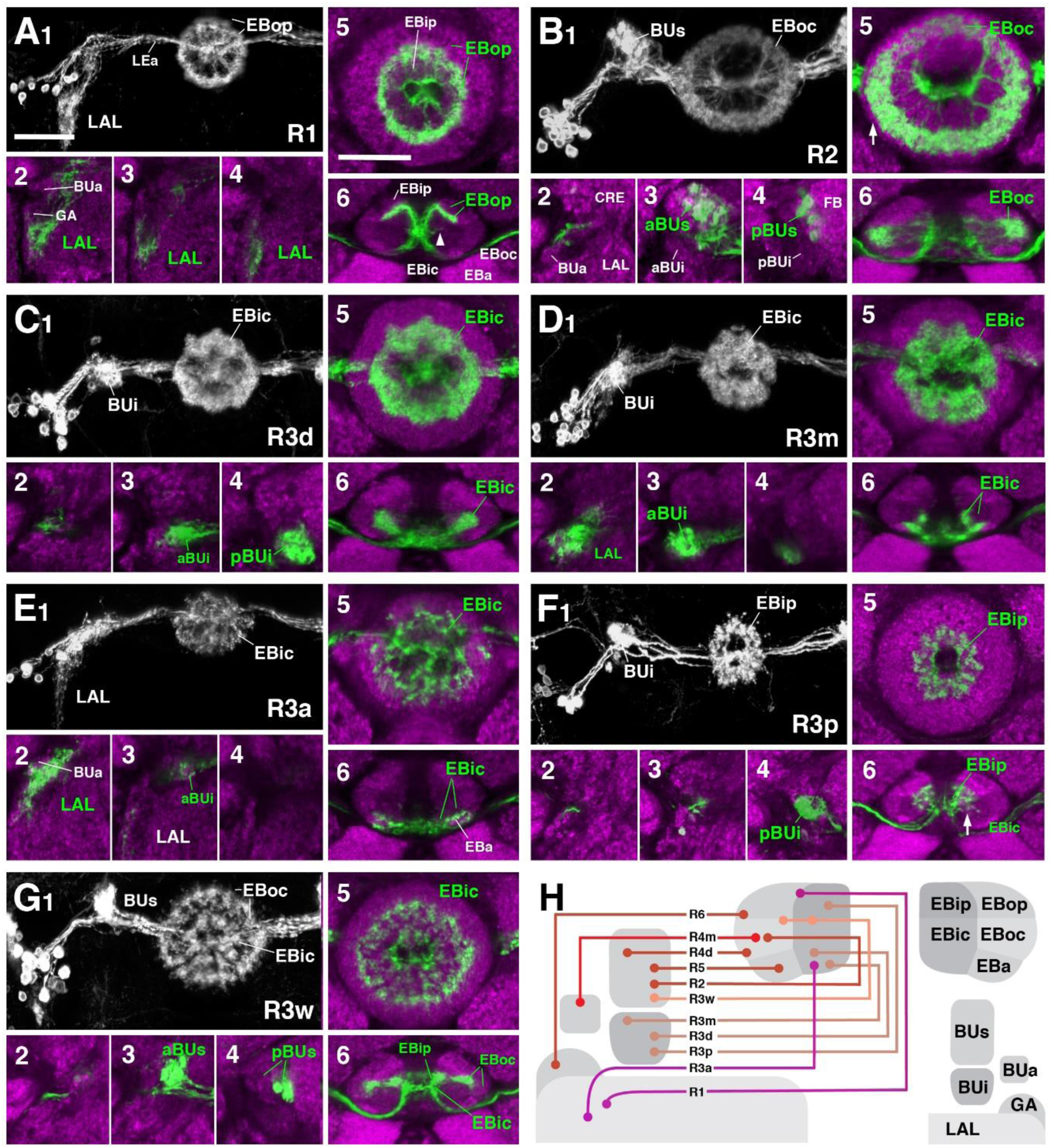
R-neuron subclasses of lineage DALv2/EBa1 with centrifugal arborizations (A-G) Confocal z-projections of Gal4 drivers that label distinct R-neuron subclasses. Each lettered, six-paneled module corresponds to an individual driver labeled with 10xUAS-mCD8::GFP. Within each module, top left (1) is a grayscale z-projection of the specific subclass (cell bodies on left), and its corresponding name in the bottom right corner. Large white annotations designate the primary domains of innervation by the subclass. In remaining module panels, the GFP-labeled neurons are shown in green; neuropil is labeled with anti-DN-cadherin (magenta). Bottom left panels (2-4) are three frontal sections of the bulb at different antero-posterior depths (as described in Fig.1); from left to right: anterior section containing BUa, intermediate section containing aBUs and aBUi, posterior section containing pBUs and pBUi. Some subclasses (R1 and R3a) innervate the lateral accessory lobe (LAL) rather than the bulb, in which case the same sections are shown at a more ventral position. Top right (5) is a higher magnification, frontal view of the ellipsoid body at an antero-posterior level (anterior, intermediate, or posterior) that highlights the circular arbor of a given driver most clearly. Bottom right (6) is a horizontal section visualizing all five DN-cadherin positive domains. Highlighted in large green text is the domain predominantly innervated by the R-neuron subclass; smaller green text signifies additional regions of innervation. Small white text in all panels denotes relevant spatial landmarks. (A1-6) R31A12-Gal4 (R1). (A1-5) Refer to Supp. Movie 1. (A1) R1 projects from the LAL to EBop. For all DALv2 R-neurons, the anterior component of the lateral ellipsoid fascicle (LEa) comprises the bridge between proximal and distal, annular neurites. (A2-4) Ventral neurites of R1 extend in the lateral LAL, medially adjacent of the gall (GA). (A5) Posterior EB section. (A6; Refer to Supp. Movie 2) R1 neurites in the EB line the anterior-most border of EBop, along the EBip-EBop interface. Additional, very small projections emanate from canal projections, along the EBip-EBic interface (arrowhead; A6). (B1-6) R78B06-Gal4 (R2). (B1-5) Refer to Supp. Movie 3. (B1) R2 projects from BUs to EBoc. (B2) No significant innervation in BUa; GFP signal corresponds to bypassing neurites. (B3) R2 neurons exhibit most of their microglomeruli in aBUs, but (B4) also some in pBUs. (B5) Intermediate EB section; arrow indicates peripheral fringe of EBoc which is not innervated. (B6; Refer to Supp. Movie 4) R2 exhibits restricted innervation of EBoc; GFP signal in EBic is passing neurites. (C1-6) R80C07-Gal4 (R3d - *distal*). (C1-5) Refer to Supp. Movie 5. (C1) R3d projects from BUi to EBic. (C2) No significant innervation in BUa; GFP signal corresponds to bypassing neurites. (C4) R3d neurons exhibit most of their microglomeruli in pBUi, but (C3) also some in aBUi. (C5) Intermediate EB section. (C6; Refer to Supp. Movie 6) R3d fills most of EBic. (D1-6) R28E01-Gal4 (R3m - *medial*). (D1-5) Refer to Supp. Movie 7. (D1) R3m projects from BUi to EBic. (D3) R3m neurons exhibit most of their microglomeruli in aBUi, but (D2) possibly also extend fibrous projections in the LAL, adjacent to BUa. (D4) No significant innervation in pBU. (D5) Anterior EB section. (D6; Refer to Supp. Movie 8) R3m fills complementary region of EBic relative to R3d. (E1-6) R12G08-Gal4 (R3a - *anterior*). (E1-5) Refer to Supp. Movie 9. (E1) R3a projects from the LAL to EBic. (E2) The LAL projections of R3a neurons are more closely related to BUa, than that of R1 neurons (A2), and (E3) may also exhibit very sparse projections in aBUi. (E4) No significant innervation in pBU. (E5) Anterior EB section. (E6; Refer to Supp. Movie 10) The EB neurites of R3a surround EBa. (F1-6) VT063949-Gal4 (R3p - *posterior*). (F1-5) Refer to Supp. Movie 11. (F1) R3p projects from BUi to EBip. (F2-3) No significant innervation in BUa and aBU; GFP signal corresponds to bypassing neurites. (F4) R3p neurons exhibit their microglomeruli in pBUi. (F5) Posterior EB section. R3p neurites in the EB densely fill EBip, but (F6; Refer to Supp. Movie 12) also appear to project anteriorly, encroaching on EBic. (G1-6) VT057232-Gal4 (R3w - *wide*). (G1-5) Refer to Supp. Movie 13. (G1) R3w projects from BUs to EBic. (G2) No significant innervation in BUa; GFP signal corresponds to bypassing neurites. (G3-4) R3w neurons exhibit their microglomeruli in aBUs and pBUs. (G5) Posterior EB section. (G6; Refer to Supp. Movie 14) R3w neurites line the posterior border of EBic, and extend into EBip. Neurites also extend distally towards EBoc, and may encroach on it. (H) Schematized overview of R-neuron domain innervation patterns (Fig.2/3 included). Other abbreviations: CRE, crepine.

**Figure 3.**
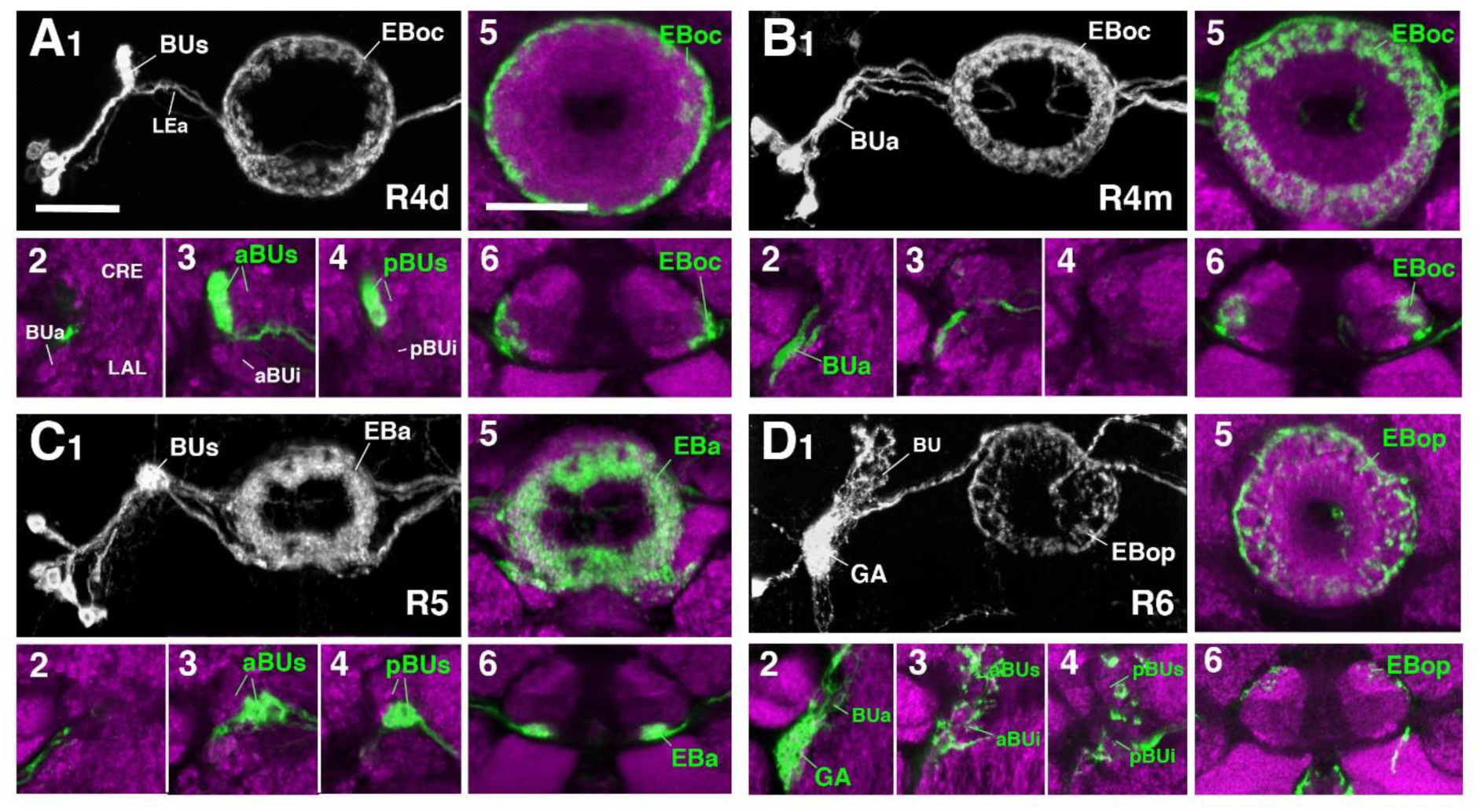
R-neuron subclasses of lineage DALv2/EBa1 with centripetal arborizations (A-D) Similar modular format of panels as described in legend for Fig. 2. (A1-6) R12B01-Gal4 (R4d - *distal*). (A1-5) Refer to Supp. Movie 15. (A1) R4d projects from BUs to EBoc. (A2) No significant innervation in BUa; GFP signal corresponds to bypassing neurites. (A3-4) R4d neurons exhibit their microglomeruli in aBUs and pBUs. (A5) Intermediate EB section. (A6; Refer to Supp. Movie 16) R4d neurites line the distal-most border of EBoc, and typically extend from posterior edge of EBa to the anterior edge of EBop. (B1-6) R59B10-Gal4 (R4m - *medial*). (B1-5) Refer to Supp. Movie 17. (B1) R4m projects from BUa to EBoc. (B2-4) Innervation only in BUa. GFP signal in aBU corresponds to bypassing neurites. (B5) Intermediate EB section. (B6; Refer to Supp. Movie 18) R4m neurites innervate EBoc. (C1-6) R58H05-Gal4 (R5). (C1-5) Refer to Supp. Movie 19. (C1) R5 projects from BUs to EBa. (C2) No significant innervation in BUa; GFP signal corresponds to bypassing neurites. (C3-4) R5 neurons exhibit their microglomeruli in aBUs and pBUs. (C5) Anterior EB section. (C6; Refer to Supp. Movie 20) R5 neurites innervate EBa. (D1-6) VT011965-Gal4 (R6). (D1-5) Refer to Supp. Movie 21. (D1) R6 projects from the Gall to EBop. (D2-4) R6 neurites are highly dense in the gall (GA), but also diffusely innervates all domains of BU. (D5) Anterior EB section. (D6; Refer to Supp. Movie 22) R6 form sparse projections in the posterior boundary of EBop and extend fine processes anteriorly into EBop. Other abbreviations: CRE, crepine; LAL, lateral accessory lobe; LEa, anterior component of the lateral ellipsoid fascicle.

### R1

R1 neurons (Renn et al., 1999), here labeled by the driver 31A12-Gal4 (Fig.2A; Supp. Movie 1 and 2; 13±2 neurons PBH), are among the minority of R-neurons that do not form glomerular dendritic branches in the bulb, but instead connect to the adjacent LAL. Here, terminal branches form a tuft of fine fibers spreading along the lateral surface of the LAL, ventrally and posteriorly adjacent to the gall (GA; Fig.2A1-A4). Distal fibers of R1 continue medially along the LE, curve around the anterior surface of the EB and, after entering the central canal, project posteriorly. Terminal distal branches densely fill a narrow volume within the outer posterior domain of the EB (EBop) which lines the boundary between EBop and the inner posterior domain (EBip) (Fig.2A5-A6). The identical pattern is labeled by c105 (Renn et al., 1999; Supp. Fig.1A). The expression of constructs that are specifically targeted towards the dendritic and axonal compartments of neurons (UAS-DenMark, UAS-syt.EGFP; Nicolaï et al., 2010) suggest that proximal projections to the LAL are exclusively postsynaptic/dendritic, and distal ones terminating in the EB are predominantly axonal (Supp. Fig.2).

### R2

R2 neurons, defined by Hanesch et al. (1989) are labeled here by 78B06-Gal4 (Fig.2B; Supp. Movie 3 and 4, 23±2 neurons PBH). Other drivers predominantly labeling this subclass are c42 (Supp. Fig.1B), introduced by Renn et al. (1999), and EB-1 (Seelig and Jayaraman, 2013; Thran et al., 2013; Young and Armstrong, 2010b; Supp. Fig.1C). R2 comprises outer R-neurons with distal, axonal endings branching throughout most of the outer central domain of the EB (EBoc; Fig.2B1, B5, B6). Only a narrow fringe along the periphery of EBoc is devoid of R2 terminals (Fig.2B5; arrow). This fringe is innervated by R4d (see below). R2 axons reach their destination by following the LE around the anterior surface of the EB, entering the central canal and then projecting centrifugally towards EBoc (Fig.2B5, B6). By this criterion they differ from the outer ring neuron subclass R4m, which projects to the same domain, but reaches it in a centripetal fashion (see below). Proximal dendrites of R2 form club-shaped glomerular endings in the medial two-thirds of the superior bulb (Fig.2B3, B4).

### R3 (distal, medial, anterior, posterior, wide)

The former R3 subclass of inner ring neurons can be broken up into at least five subclasses, R3d, R3m, R3a, R3p, R3w, defined by their axonal projections to distinct regions within the inner central (EBic) and inner posterior (EBip) domains of the EB. R3d neurons, marked by the driver 80C07-Gal4 (Fig.2C; Supp. Movie 5 and 6, 36±1 neurons PBH), project along the LE towards the anterior EB surface, turn posteriorly into the central canal and from there spread centrifugally throughout EBic (Fig.2C5, C6). They spare a small medial and anterior portion of EBic, which are innervated in a partially overlapping fashion by the subclasses R3m (Fig.2D5-D6; Supp. Movie 7 and 8, 21±1 neurons PBH) and R3a (Fig.2E5-E6; Supp. Movie 9 and10, 11±0 neurons PBH), respectively. Microglomerular dendritic endings of R3d fill the posterior region of the inferior bulb (pBUi; Fig.2C4). A few additional microglomeruli are observed in the dorso-medial part of the anterior region of the inferior bulb (aBUi), leaving the rest of aBUi empty (Fig.2C3). Dendritic projections of R3m [marked by 28E01-Gal4 in Fig.2D, and 28D01-Gal4 (Supp. Fig.1D; referred to as “R1” in Ofstad et al., 2011)] and R3a (12G08-Gal4; Fig.2E) differ from that of R3d: R3m dendrites are relatively confined to aBUi (Fig.2D3), and R3a may send very few, fibrous dendrites to aBUi, directing most of its dendrites to a small region near the dorso-lateral surface of the LAL, ventro-medially adjacent to the anterior bulb (BUa; Fig.2E1-E3). This “aberrant” dendritic projection (no glomerular synapses; targeting areas outside the bulb) puts R3a in close proximity to R1 (see above). However, the LAL territory innervated by R1 appears larger, and is located more postero-medially, than the one targeted by R3a (compare Figs.2A1-A4 and 2E1-E4).

The inner ring neuron subclass R3p, marked by VT063949-Gal4 (Fig.2F; Supp. Movie 11 and 12, 9±0 neurons PBH), has axonal projections predominantly restricted to the inner posterior domain (EBip) of the EB, largely non-overlapping with the projection of R3d/m/a (Fig.2F5-F6). Some small projections extend anteriorly, encroaching on EBic (Fig.2F6; arrow). Dendritic branches comprise a main conglomerate of endings located in the dorsal part of pBUi (Fig.2F4), a region not occupied by dendritic endings of R3d (compare to Fig.2C4).

Finally, the arborization pattern of ring neurons called R3w, marked by VT057232-Gal4 (Fig.2G; Supp. Movie 13 and 14, 27±2 neurons PBH), comes closest to that of the neuron subclass originally designated “R3” by Hanesch et al. (1989). Like all other R3 neurons listed above, R3w neurons project axons past the anterior surface of the EB into the canal, from where they spread centrifugally (Fig.2G1, G5-G6). However, terminal branches are given off centrally, near the boundary between EBic and EBip, as well as more peripherally, in the inner part of EBoc (Fig.2G5-G6). Glomerular dendritic endings occupy a medial region within the superior bulb (BUs; Fig.2G3-G4)

Driver lines expressed in inner ring neurons (“R3”) described previously may represent composites marking two or more different neuron subclasses. For example, 189y (Renn et al., 1999; Supp. Fig.1E) and 84H09 (Omoto et al., 2017; Supp. Fig.1F) include dendritic and axonal territories of R3d and R3p, targeting EBic and EBip, and much of BUi, including the dorsal territory labeled by the R3p. It is also possible that presumably composite lines 189y and 84H09-Gal4 may mark a distinct subclass of ring neurons that individually branch in both domains, which we provisionally refer to as R3c* (central). c232 and c507 (Renn et al., 1999; Supp. Fig.1G and H), as well as 15B07-Gal4 (Ofstad et al., 2011; Supp. Fig.1I) may represent a composite of R3d, R3p, and R4d, targeting EBic and EBip. The peripheral outer fringe, labeled by R4d (see below), is also present. Accordingly, most of BUi (including the dorsal R3p-associated region) and the lateral part of BUs (characteristic of R4d) are labeled by c232, c507, and 15B07-Gal4. Alternatively, these lines may represent composites of R3c* and R4d.

### R4 (distal and medial)

Two subclasses of outer ring neurons already described in the existing literature are R4d and R4m. Axons of R4d (Renn et al., 1999), here specifically labeled by 12B01-Gal4 (Fig.3A; Supp. Movie 15 and 16, 8±0 neurons PBH), reach the periphery of the EB from where they centripetally project very short terminal tufts into the fringe of EBoc. Dendritic branches form glomerular synapses confined to the lateral part of BUs (Fig.3A3-A4). R4m (Renn et al., 1999; here labeled by 59B10-Gal4) has similar axonal projections that penetrate the EB in a centripetal fashion. However, tufts of terminal branches are longer, filling the entire EBoc (Fig.3B5-B6; Supp. Movie 17 and 18, 11±3 neurons PBH) The restricted dendritic projection into BUa is highly characteristic of R4m (Fig.3B2).

### R5

In a previous paper (Omoto et al., 2017) we described an additional subclass of outer ring neurons called R5, labeled by 58H05-Gal4, which specifically targets the small, anterior EB domain (EBa; Fig.3C; Supp. Movie 19 and 20, 14±1 neurons PBH). Axons of R5 behave like those of other outer ring neuron subclasses, approaching the antero-lateral surface of the EB, and projecting short terminal branches centripetally into EBa (Fig.3C5-C6). Dendritic terminals are confined to BUs, like those of the other outer ring neuron subclasses (R2 and R4d), but occupy a small distinct locus located ventro-medially (Fig.3C3-C4). The driver 38H02-Gal4 has been described in previous works as a subset of R4 neurons (Dus et al., 2013; Ofstad et al., 2011; Park et al., 2016), but reflects a composite marker of R5 in addition to R4m (Supp. Fig.1J), with axonal projection into EBoc and EBa (Supp. Fig.1J1-J3), and dendritic endings in BUa and the ventro-medial part of BUs (Supp. Fig.1J4-J6). R5 has been referred to as “R2” or an “R2 subset” in previous works (Lin et al., 2013), particularly in the field of sleep regulation (Donlea et al., 2018; Liu et al., 2016), but is distinct from R2 as defined in classical works and herein (Hanesch et al., 1989; Renn et al., 1999).

### R6

We identified driver line VT011965-Gal4 as being expressed in a small number of DALv2 R-neurons, a subclass we refer to as R6 (Fig.3D; Supp. Movie 21 and 22, 2±0 neurons PBH). Distal neurites of these neurons approach the EB peripherally (centripetal projection) and form a sparse mesh along the posterior and postero-lateral boundary of EBop (Fig.3D1, D5-D6). They also display short branches that extend anteriorly into EBop (Fig.3D6). Proximal neurites have a unique projection pattern, first forming dense branches within the gall (GA), and then continuing into the BU, where they show a web-like innervation reaching throughout BUa, BUi, and BUs (Fig.3D2-D4).

We hypothesized that the different DALv2 subclasses are sublineages, consisting of neurons born at different time points. To address this hypothesis, we induced MARCM clones (Lee and Luo, 1999) using a pan-neuronal driver (tub-Gal4 or elav^C155^-Gal4) at defined developmental stages between 20-144h (reared at 18 °C) after hatching. Depending on where in the lineage the recombination event occurs, three types of clones [multi-cell or two-cell neuroblast clone, one-cell ganglion mother cell (GMC) clone] appear (Fig.4A). In a collection of GMC clones we found that early clone induction (20-48h) produced exclusively R4m ring neurons (Fig.4B, C). By contrast, induction at 48-72h and 72-96h resulted in a small fraction of R4m clones (approximately 20% at 48-72h, and 10% at 72-96h). Instead, we predominantly find R5, R4d, and R3 neurons (Fig.4B, D). R3 neurons subclasses are never seen in early clones (20-48h), and form the only type of clone induced at later time stages (96-144h; Fig.3B, E); 60% of the clones produced at intermediate time points belong to the R3 subclasses. These data indicate that birthdates of ring neurons differ systematically, and suggest that anatomically defined subclasses of R-neurons are indeed sublineages of DALv2.

### Posterior ExR-neurons: lineages DM4-DM6

Aside from the above described DALv2 neurons, we also observe an additional type of ring neurons. Given their widespread arborization outside the ellipsoid body, we classify them as extrinsic ring neurons (ExR), in accordance with Hanesch et al. (1989) who introduced this distinction. At least three subclasses of ExR-neurons with cell bodies located in the posterior brain cortex were recognized, and we refer to this group as the posterior ExR-neurons. Based on soma location and axonal projection these cells form part of the type II lineages CM4, CM3, and CM1, commonly known as DM4-6. Thus, all posterior ExR-neurons share a characteristic projection along the medial equatorial fascicle (MEF), which carries long axons from the posterior cortex to the LAL, extending medially of, and parallel to, the peduncle (Ito et al., 2014; Pereanu et al., 2010; Fig.5A, I, M). Fibers of large subsets of neurons belonging to lineages CM1 (DM6), CM3 (DM5), and CM4 (DM4) make up the bulk of the MEF (Ito et al., 2013; Lovick et al., 2013; Pereanu and Hartenstein, 2006; Wong et al., 2013; Yu et al., 2013).

### ExR1

The first subclass of ExR-neurons has been dubbed recently as “helicon cells” (Donlea et al., 2018), and can be visualized by the driver line 78A01-Gal4. Projecting anteriorly along the MEF, helicon axons reach the bulb and form ultra-dense arborizations in all domains of this compartment (Fig.5C). Three fiber bundles continue from the bulb towards the FB, EB, and GA of the LAL, respectively. The fiber bundle towards the FB exits BUs in the dorso-medial direction and fans out into a plexus of terminal fibers that spread along the anterior edge of the FB roof (layer 8 after Wolff et al., 2015; Fig.5D-E). The fiber bundle destined for the EB follows the LE medially. Terminal fibers form a fine web surrounding the surface of the EB. In addition, endings are concentrated in EBa and EBic (Fig.5C, E). A third contingent of fibers projects from BUa towards antero-laterally and densely innervates the GA of the LAL (Fig.5B). Based on the striking morphological similarity, helicon cells likely correspond to the first type of extrinsic R-neuron (ExR1) as defined in Hanesch et al., (1989) (Fig. 21A in Hanesch et al., 1989; Fig.12B in Young and Armstrong, 2010b; Fig. 4A in Donlea et al., 2018). Based on their cell body position and projection along the MEF, ExR1 can be attributed to the lineages DM4-DM6, but cannot be assigned to a specific lineage in the absence of further clonal analysis.

**Figure 4.**
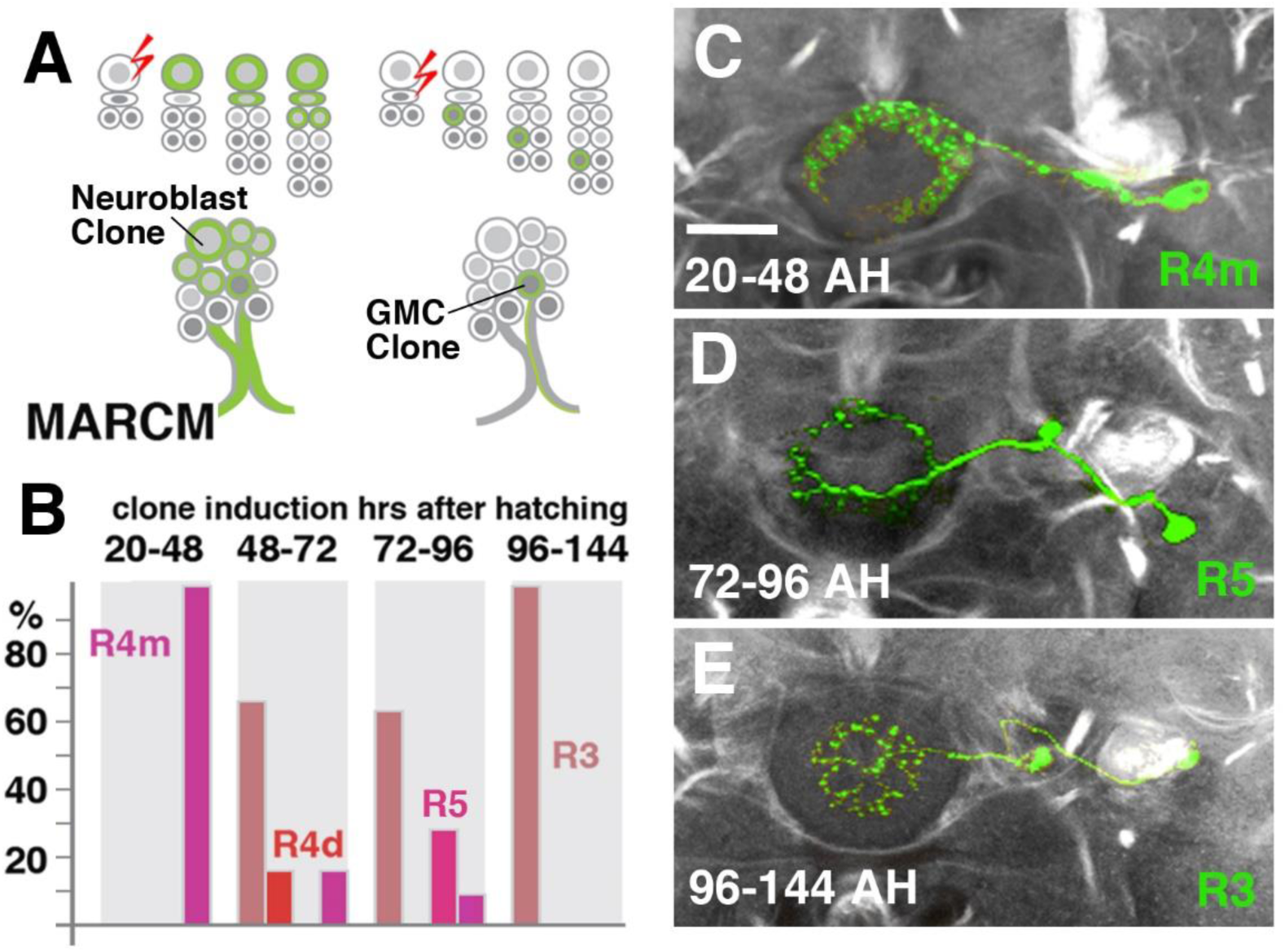
R-neuron subclasses represent sublineages of DALv2 (A) Schematized overview of MARCM clone induction and resultant clone categories. Neuroblast clones result in labeling of the entire lineage from induction onwards; ganglion mother cell (GMC) clones result in singly-labeled neurons. (B) Histogram of single cell clone R-neuron subclasses generated by temperature shifts during distinct time windows throughout larval development. A total of 33 single cell clones were generated, % reflect the proportion of clones of a given subclass generated during a given time window. (C-E) Representative clones (green) from three-time windows after hatching (AH). Axon tracts labeled by anti-Neuroglian (gray). (C) R4m (D) R5 (E) R3

### ExR2

The PPM3 group of dopaminergic neurons described in previous works encompasses the next subclass of posterior ExR-neurons (Fig.5F-I, N-Q), and can be visualized by TH-Gal4 (Friggi-Grelin et al., 2003). PPM3 is comprised of 8-9 cells whose bundled axons project anteriorly along the MEF, and form part of lineages DM4 and/or DM6 (Hartenstein et al., 2017; Ren et al., 2016). Reaching the level of the FB, PPM3 neurons branch out and innervate different compartments within the central complex (CX) and adjacent neuropils, including the bulb (BU) and lateral surface of the LAL, the anterior inferior protocerebrum [also called crepine (CRE); Ito et al., 2014], and the superior medial protocerebrum (SMP; Hartenstein et al., 2017; Fig.5F-I). Single cell clones revealed at least three different types:

**Figure 5.**
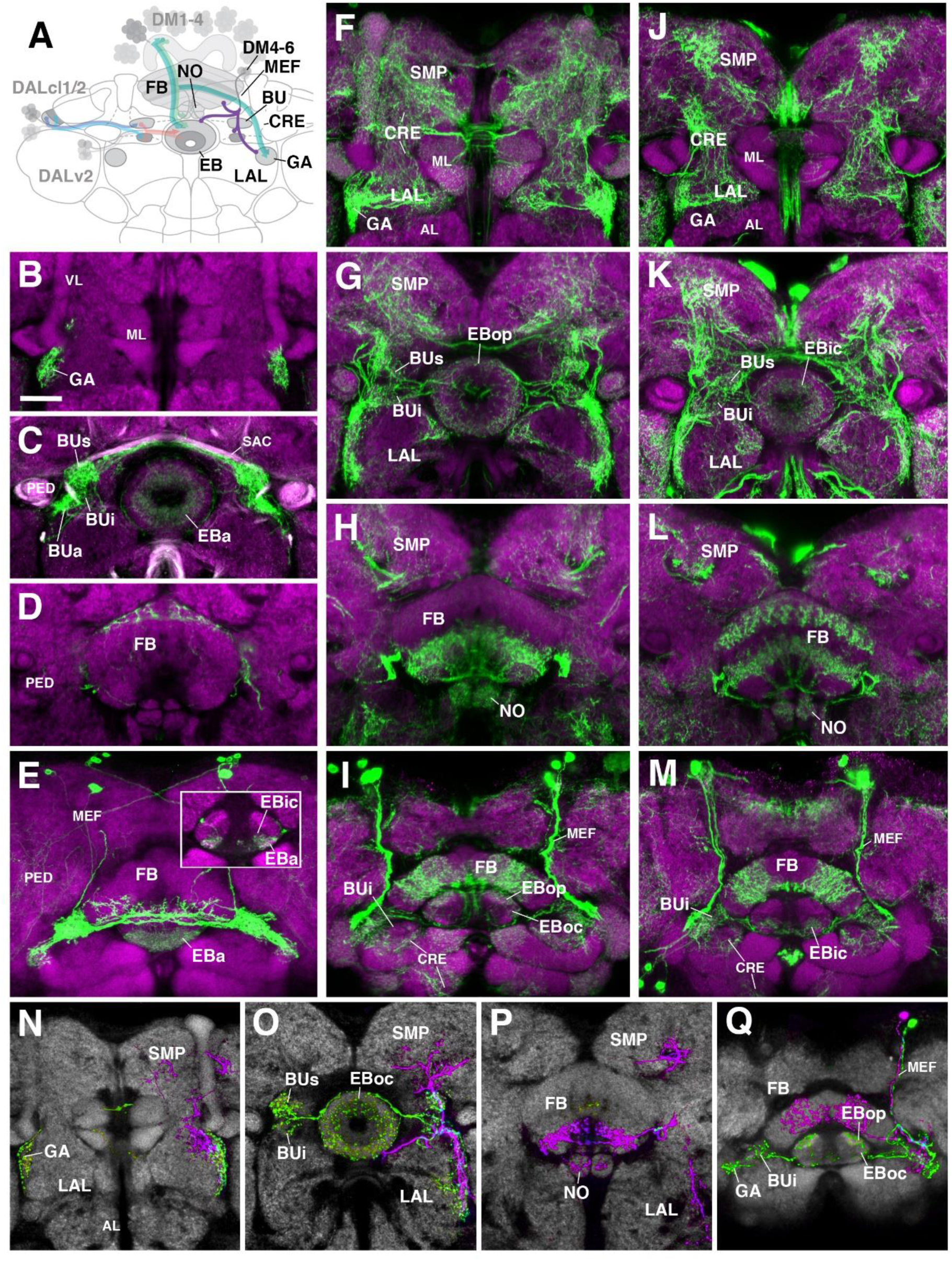
Posterior ExR-neuron subclasses of lineage CM4, CM3, CM1/DM4-6 (A) Schematized overview of interacting neuronal populations of the ellipsoid body from Fig.1, now including posterior ExR-neurons of lineages DM4-DM6 (purple). (B-M) Confocal z-projections of Gal4 drivers that label distinct ExR-neuron subclasses with posteriorly localized cell bodies. Each column corresponds to a single driver (green) labeled with 10xUAS-mCD8::GFP. Neuropil labeled by anti-DN-cadherin (magenta). Top three rows of each column correspond to frontal sections at three different antero-posterior depths; from top to bottom: anterior section containing the Gall (GA)/lateral accessory lobe (LAL), intermediate section containing the ellipsoid body (EB) and bulbs (BU), posterior section containing the fan-shaped body (FB) and noduli (NO). Bottom (fourth) row is a horizontal section visualizing the length of the EB canal. Larger white annotations denote arborization-containing domains of interest; smaller white annotations represent spatial landmarks. (B-E) R78A01-Gal4 (ExR1; “helicon cells”). Inset in (E) depicts dorsal view of the EB; ExR1 innervates EBa and the anterior part of EBic. (F-I) TH-Gal4 (ExR2 and other TH-positive neurons). (I) Dorsal view of the EB shows innervation in EBop; sparse innervation in EBoc also detected. (J-M) TPH-Gal4 (ExR3 and other TPH-positive neurons). (M) Dorsal view of the EB shows innervation in anterior part of EBic. (N-Q) Confocal z-projections of individually-labeled cells generated by MCFO using TH-Gal4. Four panels depict the same sections and are organized in the same fashion as in B-M, but arranged from left to right rather than top to bottom. Neuropil labeled by anti-DN-cadherin (gray). (N-O) Green cell is ExR2 (PPM3-EB dopaminergic neuron); a single PPM3-EB innervates the Gall (GA) of the lateral accessory lobe (LAL), lateral region of the LAL, and all BU domains on both sides. (Q) Additional innervation includes prominent infiltration into EBop, and sparser innervation of EBoc. Magenta cell is not an ExR-neuron (PPM3-FB dopaminergic neuron). Annotation format is identical to that of B-M. Other abbreviations: AL, antennal lobe; CRE, crepine; MEF, medial equatorial fascicle; ML, VL, and PED, medial lobe, vertical lobe, and peduncle of the mushroom body, respectively; SMP, superior medial protocerebrum; BUs and BUi, superior and inferior domains of the bulb; anterior (EBa), inner central (EBic), outer central (EBoc), and outer posterior (EBop) domains of the ellipsoid body.

(1) PPM3 neurons innervating the EB, BU, and lateral surface of the LAL (including the GA; Fig.5N; called PPM3-EB in the following). The PPM3-EB axon projects along the LE and enters EB at a dorso-lateral position (Fig.5O). Terminal arborizations are concentrated in EBop (Fig.5Q). Additional branches reach superior and inferior parts of both ipsi- and contralateral bulbs (Fig.5O). Note that within the BU, unlike most DALv2 ring neurons (see Figs.2 and 3) or the afferent TuBu neurons (Omoto et al., 2017), terminal PPM3-EB branches do not end in large microglomerular structures, but form thin, highly branched endings. Due to its EB innervation, we refer to these neurons as the ExR2 subclass. Like PPM3-EB, ExR2 as defined by Hanesch et al., (1989) also contains a caudal innervation pattern in the EB, but a direct correspondence cannot be made since the cell from this study was not fully reconstructed and the ring neuron from lineage BAmv1 also innervates the same EB region (see below).

(2) At least two subtly different kinds of PPM3 neurons innervating FB, NO, LAL, CRE, and SMP exist, one of them shown in Fig.5N-Q (magenta; PPM3-FB). This cell arborizes in the ventral layers (2-3, after Wolff et al., 2015) of the FB, and the intermediate noduli (NO2). Separate branches project to the lateral surface of the LAL, where projections partially overlap with those of PPM3-EB (green; Fig.5O), but stay out of the GA and instead reach the medially adjacent CRE (Fig.5N, O). A third branch projects upward into a discrete subdomain of the SMP (Fig.5O, P).

(3) A third type of PPM3 neuron (PPM3-LAL; not shown) does not innervate the EB or FB, but has bilateral projections to the lateral surface of the LAL. The second and third type of PPM3 neurons are not considered ExR-neurons due to their lack of EB innervation.

### ExR3

A subset of serotonergic neurons, visualized by the driver TPH-Gal4 (Park et al., 2006), also form part of the type II lineages, DM4-6, and have widespread projections towards the CX, BU, LAL, and CRE (Fig.5J-M). Single cell labeling to resolve cell types with different projections within the CX have not been carried out. A plexus of thin fibers enters the EB from laterally via the LE. Within the EB, terminal arborizations are largely non-overlapping with those of dopaminergic PPM3-EB (see below), showing highest density in EBic (Fig.5M). The serotonergic neuron subclass that innervates this domain is designated herein as ExR3 and is derived from DM4-6, but like ExR1, cannot be assigned to a specific lineage. Additionally, in the FB, one can distinguish innervation of a ventral stratum (layers 3-4 of Wolff et al., 2015) from that of a dorsal stratum (layers 6/7).

### Anterior ExR-neurons: lineage BAmv1/LALv1

One additional subclass of ExR-neurons, designated as ExR4, was identified. Cell bodies of these neurons form a cluster in the anterior brain cortex, but, in contrast to DALv2 R-neurons, are located ventrally of the antennal lobe (Fig.6). ExR4 neurons, labeled by the driver 14G09-Gal4 (Fig.6; Supp. Movie 23 and 24), belong to the lineage BAmv1 (Lovick et al., 2013; Wong et al., 2013), also called LALv1 (Ito et al., 2013; Yu et al., 2013).

ExR4 axons follow a highly characteristic pathway that initially leads posteriorly as part of the longitudinal ventro-medial fascicle (loVM) and then makes a sharp turn dorsally (Fig.6A, B). The dorsal leg of the BAmv1 tract penetrates the LAL and gives off dense tufts of branches that fill the dorso-lateral quadrant of the LAL compartment (Fig.6B, E). Some branches reach forward into the GA of the LAL (Fig.6B, D); others continue further dorsal into the CRE compartment (Fig.6B-D). Reaching the dorsal edge of the LAL, the axon tract of BAmv1 makes a second sharp turn, projecting medially towards the CX as the posterior part of the lateral ellipsoid fascicle (LEp). Axons reach the CX at the cleft between the EB and FB, and from there project anteriorly into EBop (Fig.6A, B, G) and posteriorly towards the ventral strata (1-4, after Wolff et al., 2015) of the FB and into NO2 (Fig.6A, B, F).

**Figure 6.**
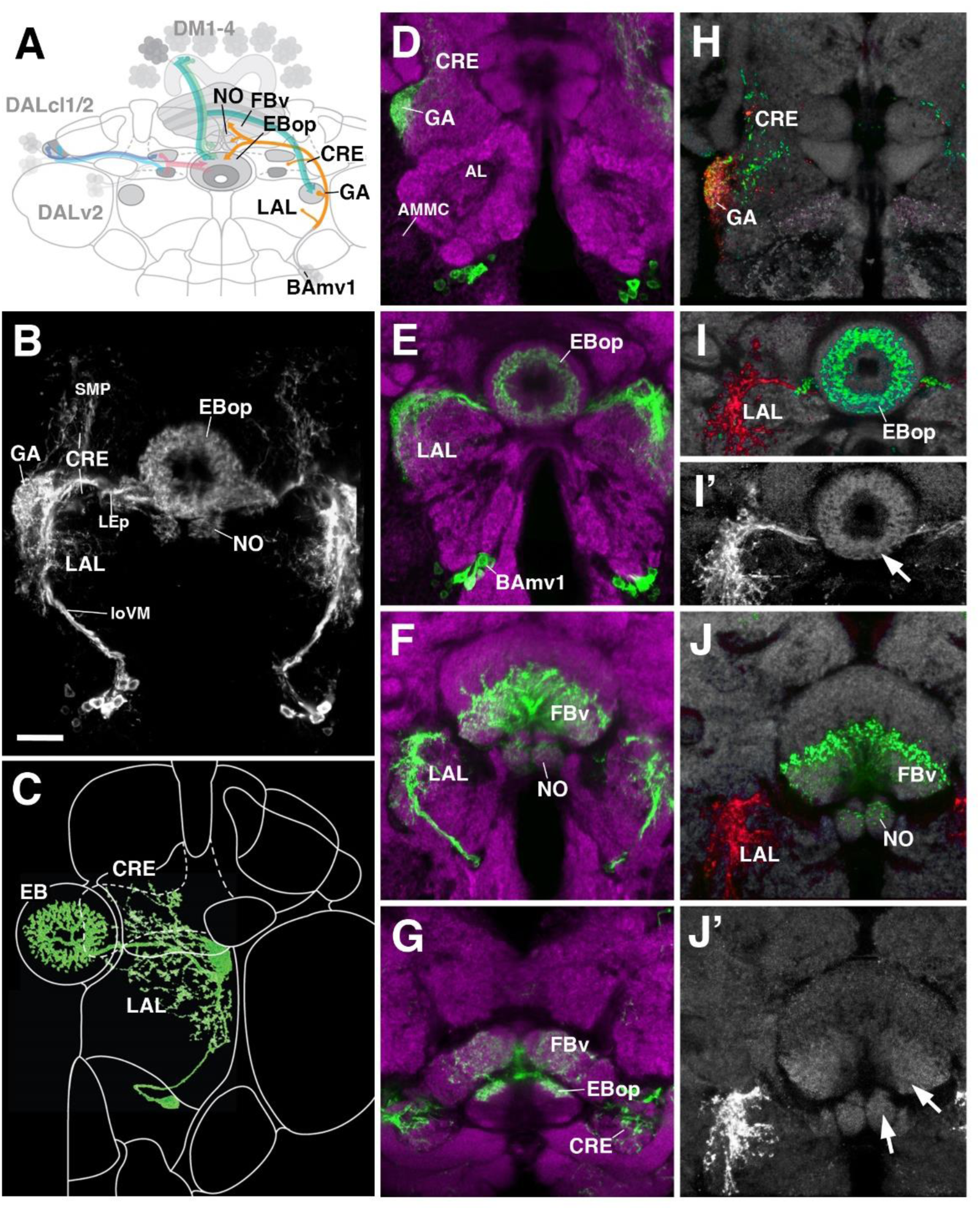
Anterior ExR-neuron subclass of lineage BAmv1/LALv1 (A) Schematized overview of interacting neuronal populations of the ellipsoid body from Fig.1, now including anterior ExR-neurons of lineage BAmv1 (orange). (B) Grayscale z-projection of ExR4, labeled by R14G09 > 10xUAS-mCD8::GFP. Z-projection spans from the Gall (GA)/lateral accessory lobe (LAL) to the EB, and does not include the fan-shaped body (FB) component of this driver (see below). ExR4 projects from the GA/LAL to EBop. (C) Single cell clone of ExR4; identified from the Flycircuit database (Chiang et al., 2011). (D-G) Top three rows of middle column corresponds to frontal sections of R14G09 > 10xUAS-mCD8::GFP at three different antero-posterior depths (Refer to Supp. Movie 23). From top to bottom: (D) anterior section containing the GA/LAL, (E) intermediate section containing the EB and bulbs, (F) posterior section containing the FB and noduli. Bottom row (G; Refer to Supp. Movie 24) is a horizontal section visualizing the length of the EB canal. R14G09-positive neurons are shown in shown in green; neuropil is labeled with anti-DN-cadherin (magenta). (H-J) Corresponding sections in D-F using R14G09 labeled with the presynaptic marker syt.EGFP (green) and dendritic marker DenMark (red). (H, I, J) Neuropil is labeled with anti-DN-cadherin (gray). I’ and J’ show the isolated DenMark signal from I and J in white. Other abbreviations: AL, antennal lobe; AMMC, antenno-mechanosensory and motor center; CRE, crepine; FBv, ventral region of the fan-shaped body; LEp, posterior component of the lateral ellipsoid fascicle; loVM, medial component of the ventral longitudinal fascicle; SMP, superior medial protocerebrum.

The structure of individual ExR-neurons of the BAmv1 lineage have not been thus far described in the literature. We identified a single cell clone in the FlyCircuit database (Chiang et al., 2011) that, based on axonal trajectory (loVM, LEp, EB), represents BAmv1 (Fig.6C). Notably, this neuron has profuse branches in the LAL (including the GA) and in the posterior EB, but does not project to the FB or NO, indicating that different neuron types of the BAmv1 lineage innervate the EB and the FB/NO, respectively.

DenMark and syt.EGFP expression revealed that projections of BAmv1 neurons towards the EB and FB/NO are mainly axonal, with only a weak dendritic component; proximal arborizations in the LAL are preferentially dendritic (Fig.6I-J’). Only the gall of the LAL has a significant axonal component (Fig.6H). This distribution of pre- and postsynaptic elements suggests that ExR-neurons of BAmv1 form a feed-back component connecting input and output domains within the CX circuitry: DALv2 ring neurons provide strong input to the EB, which is then transmitted to the FB and the GA/LAL by columnar neurons. BAmv1 neurons form dendritic endings in the GA/LAL and feedback axons towards the EB.

### Columnar neurons

While ring neurons terminate in the EB, further processing of visual input requires synaptic partners that access other compartments of the CX. Several populations of columnar neurons fulfilling this requirement have been identified to date (Wolff et al., 2015). To investigate the spatial relationship of columnar neurons with ring neurons, we screened Gal4 drivers and identified lines with distinct columnar expression patterns (Fig.7). In this study, we anatomically assess four of such populations. These are two populations of “wedge neurons” with arborizations in both peripheral and central parts of the EB, and two populations of “tile” neurons that project only to the peripheral EB. Shown in Fig.7A is the first type of “wedge” neuron (PB_G1–8_.b-EBw.s-D/Vgall.b), also called E-PG neurons (Turner-Evans et al., 2017; Wolff et al., 2015), whose spiny (dendritic) terminal branches fill all EB domains except for much of EBa (Fig.7A1-A2). Confirming previous descriptions, bulbar (axonal) endings are seen in the PB, as well as the GA (Fig.7A1, A3, A6).

**Figure 7.**
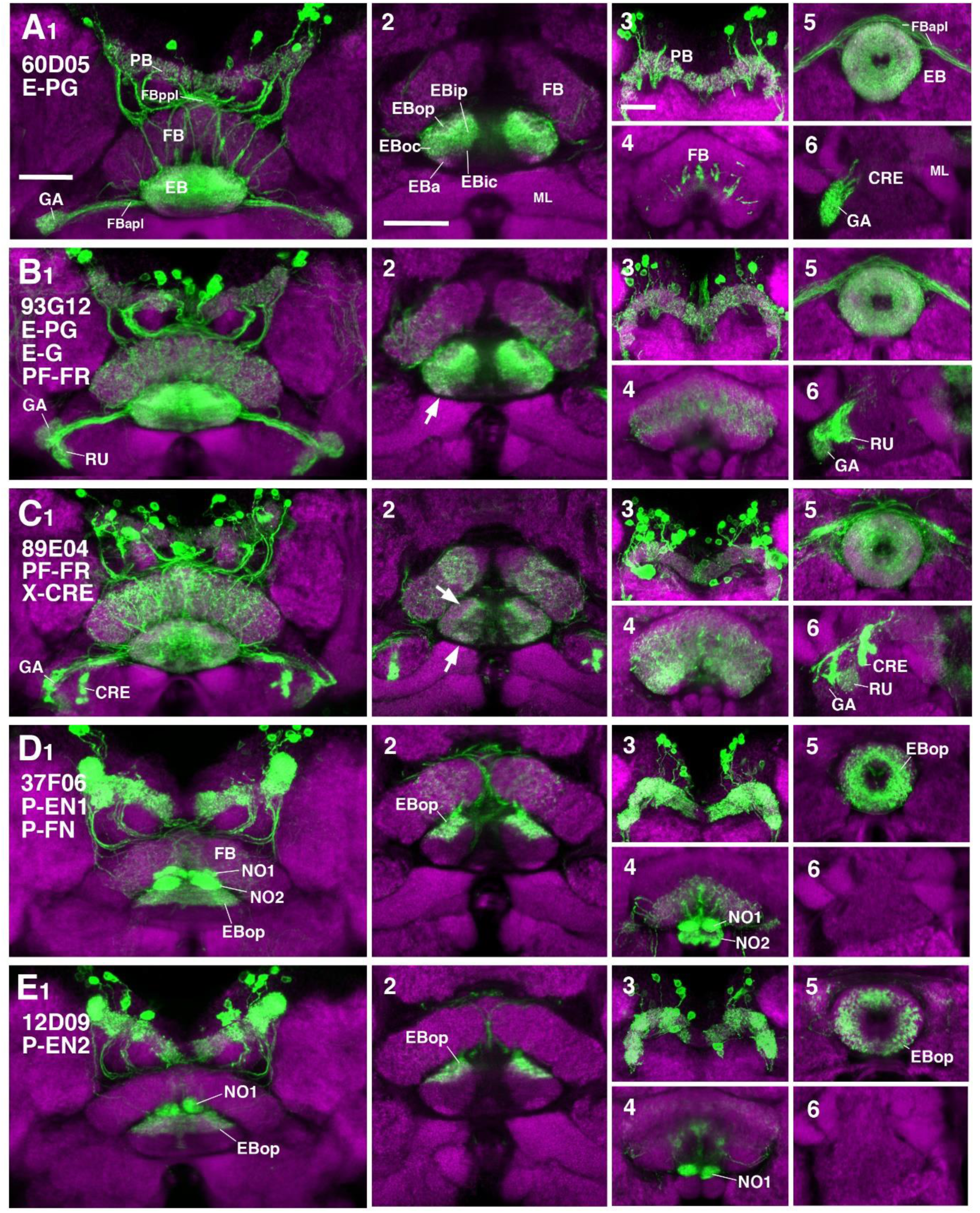
CX innervation patterns of Gal4 drivers labeling columnar neuron populations (A-E) Confocal z-projections of Gal4 drivers that label columnar neuron classes. Each lettered, six-paneled module corresponds to an individual driver labeled with 10xUAS-mCD8::GFP (green); neuropil is labeled with anti-DN-cadherin (magenta). Left (1) is a z-projection from the dorsal view spanning the antero-posterior depth of the brain (anterior pointing downward). Second column (2) is a horizontal section visualizing the length of the EB canal; all five DN-cadherin positive domains are visible. Final two columns (3-6) are four frontal sections of the CX and its associated neuropils: (3) protocerebral bridge (PB), (4) fan-shaped body (FB), (5) ellipsoid body (EB), (6) gall (GA)/lateral accessory lobe (LAL). Larger white annotations denote arborization-containing domains of interest; smaller white annotations represent spatial landmarks. (A1-6) R60D05-Gal4 (E-PG neurons). (B1-6) R89E04-Gal4 (E-PG, “E-G”, “PF-FR” neurons). (2) Similar distribution of GFP signal as E-PG neurons (A2), with additional innervation of EBa (arrow). (C1-6) R93G12-Gal4 [“PF-FR” neurons, “X-CRE” indicates a novel neuronal population with glomerular endings in the crepine (CRE), dendritic terminals and their domains of innervation can not be resolved]. (2) Ubiquitous innervation of the EB, with enriched signal in EBa and the anterior border of EBop (arrows). (D1-6) R37F06-Gal4 (P-EN1, “P-FN” neurons). (2) EB innervation is restricted to outer posterior domain (EBop). (E1-6) R12D09-Gal4 (P-EN2). (2) EB innervation is restricted to outer posterior domain (EBop). Other abbreviations: FBapl and FBppl, anterior and posterior plexus of the fan-shaped body; RU, rubus.

The second type of “wedge” neuron described to date (EBw.AMP.s-Dga-s.b; Wolff et al., 2015) has dendritic endings confined to the EB and axonal projections to a region adjacent to the dorsal gall, called the dorsal gall surround (Dga-s). For brevity, we provisionally refer to this neuronal population as “E-G” neurons. In the EB, terminal arbors fill all domains, including EBa. Driver line 93G12-Gal4 (Fig.7B) is likely to include this neuron type, given strong label of the entire EB (including EBa) and the Dga-s. However, other neuron populations are marked as well, including, most likely, PB_G1–8_.s-FBl3,4,5.s.b-rub.b (provisionally “PF-FR”) neurons [label in the PB, FB, and rubus (RU) domain of the CRE] and E-PG neurons (label in the PB and ventral GA) (Wolff et al., 2015; Fig.7B).

The list of columnar neurons described in the literature is likely to be incomplete, given that the comprehensive overview of Wolff et al. (2015) focused primarily on neurons with connections to the PB. Another such cell type, not yet described, is included in the population of neurons labeled by 89E04-Gal4 (Fig.7C). Enriched signal is seen in EBip and EBa (Fig.7C1, C2; arrows), a pattern not shown by E-PG neurons or P-EN neurons. Presumed endings of this cell type form dense glomerular structures in the CRE (Fig.7C1, C2, C6) and the Dga-s. As the compartments which contain dendritic components of these neurons cannot be precisely resolved, we provisionally refer to them as “X-CRE” neurons. PF-FR neurons are likely marked by 89E04-Gal4 as well, given the label in these compartments (Fig.7C3, C4, C6).

Fig.7D, E show the projection pattern of P-EN neurons (PB_G2–9_.s-EBt.b-NO_1_.b; Turner-Evans et al., 2017; Wolff et al., 2015). This neuronal population has axonal endings in a tile-shaped domain, which we show here corresponds to EBop (Fig.7D2, D5, E2, E5). Outside the EB, projections are in the PB (Fig.7D3, E3) and the dorsal noduli (NO1; Fig.7D4, E4). Functionally, P-EN neurons fall into two subclasses, one (called P-EN1; Green et al., 2017; Turner-Evans et al., 2017) marked by the driver 37F06-Gal4 (Fig.7D), and the other one (P-EN2; Green et al., 2017) by 12D09-Gal4 (Fig.7E). In regard to terminal arborization in the EB, both lines appear identical (compare Fig.7D2, E2). However, 37F06-Gal4 is likely expressed in an additional neuron type called PB_G2–9_.s-FBl3.b-NO_2_V.b (provisionally “P-F_3_N_2_”; subscripts denote which subcompartment of the FB or NO is innervated), given labeling in the FB and NO2 (Wolff et al., 2015; Fig.7D4).

### Mapping the putative postsynaptic targets of R-neurons

To identify downstream postsynaptic targets of distinct R-neuron subclasses, we utilized the anterograde trans-synaptic tracing method *trans*-Tango (Talay et al., 2017). In this approach, a bioengineered synthetic receptor system and downstream signaling components are expressed in a pan-neuronal fashion. Gal4-dependent expression of a presynaptically-tethered cognate ligand leads to activation of the receptor specifically in postsynaptic neurons downstream of the Gal4 expressing population. Receptor activation then leads to proteolytic cleavage and release of the otherwise membrane-sequestered, orthogonal transcriptional activator QF. In this manner, the presynaptic neurons labeled by GFP under UAS control, can be visualized in conjunction with downstream targets, labeled by RFP under QUAS control. We applied *trans*-Tango to every R-neuron Gal4 driver to reveal, in principle, each output system in its entirety. However, in the absence of parallel methodology to demonstrate functional connectivity, we consider our findings to reveal “putative” targets, due to several potential caveats that merit consideration, some of which have been previously noted (Talay et al., 2017). In some cases, such as in R4d and R6, *trans*-Tango did not successfully yield RFP signal in the EB. This may be a consequence of insufficient expression levels of the synthetic ligand due to a weak Gal4 line, or a neural circuit that simply contains fewer synapses. Indeed, the Gal4 drivers that label R4d and R6 appear weaker, and are also some of the smallest populations in terms of cell number (8±0 and 2±0 neurons PBH, respectively). However, a large or strongly Gal4-expressing GFP-positive population is not required to elicit a strong *trans*-Tango signal. We anecdotally observed strong RFP expression in other areas of the brain due to non-specific expression outside of the CX, even when the non-specific cells were sparse or weakly labeled. The strength of *trans*-Tango signal may also be determined by the neuron-specific expression level of the synthetic receptor, which cannot be assumed a priori to be uniformly expressed in every neuron throughout the brain, despite being under the control of a pan-neuronal promoter. This may result in sparse presynaptic labeling leading to strong postsynaptic labeling, or vice versa. This potential confound leads to the next consideration; a highly specific driver line is important to accurately interpret the postsynaptic signal. Particularly for the CX, where neurons between neuropil compartments are recurrently connected, any non-specific neuronal labeling in another CX neuropil outside of the EB may yield a false positive signal for a ring neuron driver *trans*-Tango experiment. In a hypothetical example, it would be difficult to disambiguate whether RFP-positive neurons that interconnect the PB and EB are downstream of the ring neuron class of interest, or a non-specific neuron in the PB. Considering this, we opted not to include *trans*-Tango results from R5 (58H05-Gal4); this driver line included several other neurons in other CX neuropils (data not shown). Finally, the potential for false-positive signal due to reporter sensitivity or mislocalization of the overexpressed ligand, must always be considered (Talay et al., 2017). Despite the requirement for further validation, the *trans*-Tango results herein reveal heretofore unknown wiring principles of the R-neuron network. The findings of these experiments are summarized in Fig.10.

### R1

Neurons trans-synaptically labeled when using the R1 driver 31A12-Gal4 included R-neurons of the R1 and R3p subclass, as well as columnar E-PG and, likely, PB_G2–9_.s-FBl2.b-NO_3_A.b (provisionally “P-F_2_N_3_”) neurons (Wolff et al., 2015). Thus, *trans*-Tango signal is detected in cell bodies of DALv2 neurons in the anterior cortex, as well as DM1-4 neurons in the posterior cortex (Fig.8A1, 2, 3). Many of the labeled DALv2 neurons were also positive for GFP (Fig.8A3), indicating that R1 neurons form strong reciprocal connections among each other (homotypic interactions). In the EB neuropil, *trans*-Tango signal fills all compartments, but is enriched in EBip (Fig.8A4, A10), which is targeted by R3p neurons (heterotypic interactions). Accordingly, labeling is also detected in a subset of glomeruli of BUi (Fig.8A5), the dendritic compartment of R3p. *Trans*-Tango-positive projections of columnar neurons accounts for the labeling detected in the PB (Fig.8A8), outer EB domains (Fig.8A4, A10), and GA (Fig.8A11). Signal ventrally of the GA (Fig.8A11), filling the lateral surface of the LAL, is attributable to the reciprocally connected R1 neurons (Fig.8A5-A7, A11). In addition to E-PG neurons, the *trans*-Tango-positive columnar neurons also appeared to include P-F_2_N_3_ neurons, based on signal detectable in the fan-shaped body and ventral noduli (Fig.8A9). The responsible connection between R1 and P-F_2_N_3_ neurons to which this label could be attributable, could be a sparse, posteriorly projecting neurite of R1-neurons to the FB which was periodically observed (data not shown).

**Figure 8.**
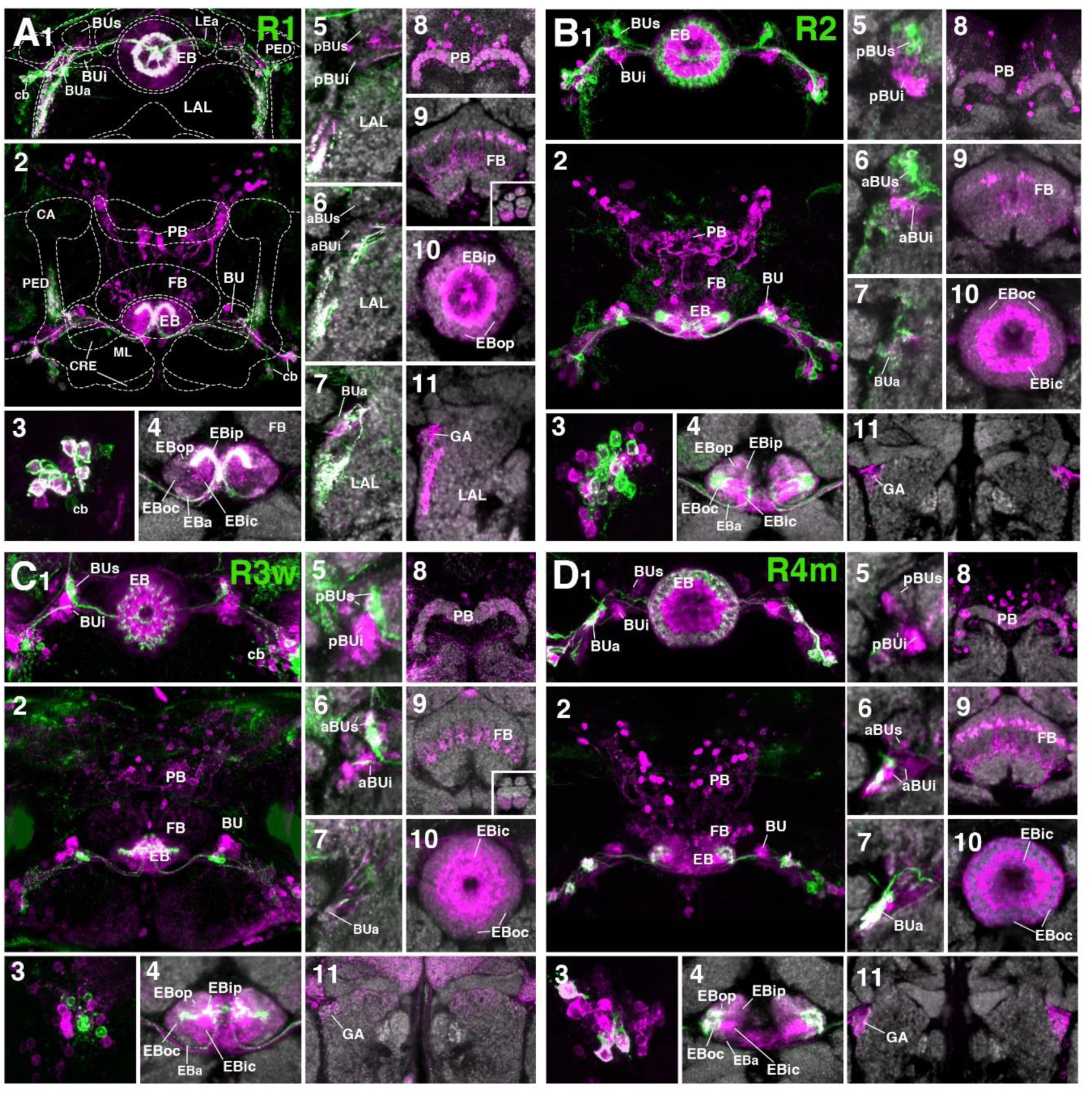
Putative postsynaptic partners of R-neurons revealed by *trans*-Tango (A-D) Confocal z-projections of Gal4 drivers that label distinct R-neuron subclasses in conjunction with *trans*-Tango mediated labeling of postsynaptic neurons. Each lettered, eleven-paneled module corresponds to an individual driver. Gal4-expressing R-neurons labeled by GFP under UAS control (green), putative postsynaptic neurons are labeled by RFP under *trans*-Tango mediated QUAS control (magenta). Larger white annotations denote arborization-containing domains of interest; smaller white annotations represent spatial landmarks. Module organization outlined below: (1) Frontal z-projection spanning from R-neuron cell bodies to the ellipsoid body (EB)/bulbs (BU). (2) Z-projection from the dorsal view spanning the antero-posterior depth of the brain (anterior pointing downward). (3) Cell body image to illustrate degree of colocalization between pre- and post-synaptic neurons, an indication of homotypic interactions within a cell type. (4-11) Neuropil is labeled with anti-DN-cadherin (gray). (4) Horizontal section of the EB spanning the length of the EB canal. (5-7) Frontal sections of the bulb at three different depths. From top to bottom: (5) posterior section containing the posterior regions of the superior (pBUs) and inferior (pBUi) bulb, (6) intermediate section containing the anterior regions of the superior (aBUs) and inferior (aBUi) bulb, (7) anterior section containing the anterior (BUa) bulb. (8-11) Isolated postsynaptic targets (shown in magenta) throughout the CX (Gal4-expressing neurons are shown not shown). Four frontal sections of the CX and its associated neuropils; from top (posterior-most) to bottom (anterior-most): (8) protocerebral bridge, (9) fan-shaped body, (10) ellipsoid body, (11) gall (GA)/lateral accessory lobe (LAL). (A1-11) (R1) R31A12-Gal4 > *trans*-Tango (B1-11) (R2) R72B06-Gal4 > *trans*-Tango (C1-11) (R3w) VT057232-Gal4 > *trans*-Tango (D1-11) (R4m) R59B10-Gal4 > *trans*-Tango Other abbreviations: CRE, crepine; ML, PED, and CA, medial lobe, peduncle, and calyx of the mushroom body, respectively; BUa, BUs, and BUi, anterior, superior, and inferior domains of the bulb, respectively; anterior (EBa), inner central (EBic), outer central (EBoc), inner posterior (EBip), and outer posterior (EBop) domains of the ellipsoid body; LEa, anterior component of the lateral ellipsoid fascicle; cb, cell bodies.

### R2

Cells identifiable as targets of R2-neurons include other ring neurons of the R4d, R3d, and R2 subclasses, in addition to columnar E-PG neurons. In the anterior cortex, GFP and *trans*-Tango label overlaps weakly, and perhaps in only a small proportion of cell bodies (Fig.8B3), indicating that reciprocal connections among R2-neurons are present but not very pronounced. The EB neuropil is ubiquitously filled with trans-synaptic label, but shows enriched signal in EBic, which is targeted by R3d, and the outer rim of EBoc, innervated by R4d (Fig.8B4), suggesting prominent heterotypic interactions. In accordance with the notion that R3d and R4d are postsynaptic partners of R2, we find *trans*-Tango labeling in a posterior-lateral part of BUs (Fig.8B5), shown above to be dendritically innervated by R4d (see Fig.3A3-A4), and in the posterior region of the inferior bulb (pBUi), corresponding to R3d (Fig.8B5). Labeling of cell bodies in the posterior cortex, occupied by DM1-4 (Fig.8B2), and of neuropil including the PB (Fig.8B8), outer EB (Fig.8B10) and GA (Fig.8B11) is attributable to E-PG neurons targeted by R2. Sparse label in the FB corresponds to the through-going fibers of columnar E-PG neurons, in addition to other neuronal populations of unclear identity (Fig.8B9).

### R3w

The R3w subclass could be viewed as an intermediary between outer and inner R-neurons in terms of the projection of proximal branches (ventral part of BUs) and distal branches (narrow EBic domain, reaching into EBoc). Correspondingly, R3w neurons, like outer subclasses R2 and R4m, target a good number of columnar E-PG neurons (Fig.8C2, C8, C11), in addition to a small number of R2 neurons (sparse RFP label in EBoc and BUs; Fig.8C4-C6). Strongest RFP signal is seen in the inner EB domains and BUi, indicating R3d, R3m, and particularly R3p, as major targets (Fig.8C4-C6, C10). It should be noted that the driver VT057232-Gal4, which is used here to label R3w, is also expressed in numerous additional neurons throughout the brain, which may account for some of the RFP signal (Fig.8C2). For example, staining of the ventral noduli (NO3; Fig.8C9, inset), presumably corresponding to one of the P-FN populations, is unlikely to be attributable to R3w R-neurons as presynaptic partners.

### R4m

R4m neurons innervate EBoc, similar to R2, but have their dendritic projection towards BUa. Using the R4m driver line 59B10-Gal4, we see trans-synaptic label of R-neuron subclasses R4m, R4d and R3d, as well as a large fraction of columnar E-PG neurons (Fig.8D). Signal is concentrated in EBic (R3d) and periphery of EBoc (R4d) (Fig.8D4, D10). In the BU, trans-synaptic label covers BUa, corroborating reciprocal interactions among R4m (Fig.8D7), in addition to the lateral part of BUs (R4d; Fig.8D5) and pBUi (R3d; Fig.8D5). There is a large number of RFP-positive DM1-4 columnar neurons in the posterior cortex (Fig.8D2), projecting dense arrays of fiber tracts that innervate the PB (Fig.8D8), through the FB (Fig.8D9), to the EB and GA (Fig.8D10-D11).

### R3d/4d

The driver line 80C07-Gal4 marks R3d neurons with axonal projections to EBic and dendritic innervation of pBUi. This pattern is confirmed when using the line in the context of trans-synaptic labeling (Fig.9A1, A5). However, likely as a consequence of enhancement of GFP signal with the use of anti-GFP antibody, we also saw GFP signal in the outer rim of EBoc and the lateral part of BUs, indicating that 80C07-Gal4 is also expressed in R4d neurons. It is therefore difficult to assign the observed trans-synaptic label induced by this line to either one of these populations. We observe trans-synaptic label in both R-neurons in the anterior cortex, and DM1-4 columnar neurons in the posterior cortex (Fig.9A1-A2). Approximately half of 80C07-expressing R-neurons are also labeled trans-synaptically, suggesting strong reciprocal connectivity among R3d and/or R4d (Fig.9A3). In the EB neuropil, *trans*-Tango signal is highly concentrated in EBic and EBip, and the outer rim of EBoc (Fig.9A4, A10). This pattern again argues for strong reciprocal interactions of R3d and R4d, as well as contacts between R3d and R3p. Trans-synaptic label in the BU also largely overlaps with GFP signal in pBUi and lateral part of BUs (Fig.9A5-A6), that are dendritically innervated by R3d and R4d neurons, respectively. Exclusive label of glomeruli in the dorsal part of BUi (Fig.9A5) corresponds to R3p which targets this region (see Fig.2F). Trans-synaptic labeling of columnar neurons is sparse, with only few cell bodies in the posterior cortex (Fig.9A2, A8), and faint/restricted labeling of the outer EB domains and the GA (Fig.9A10-A11). The identity of trans-synaptically labeled neurons spottily innervating the FB (Fig.9A9) is not clear.

**Figure 9.**
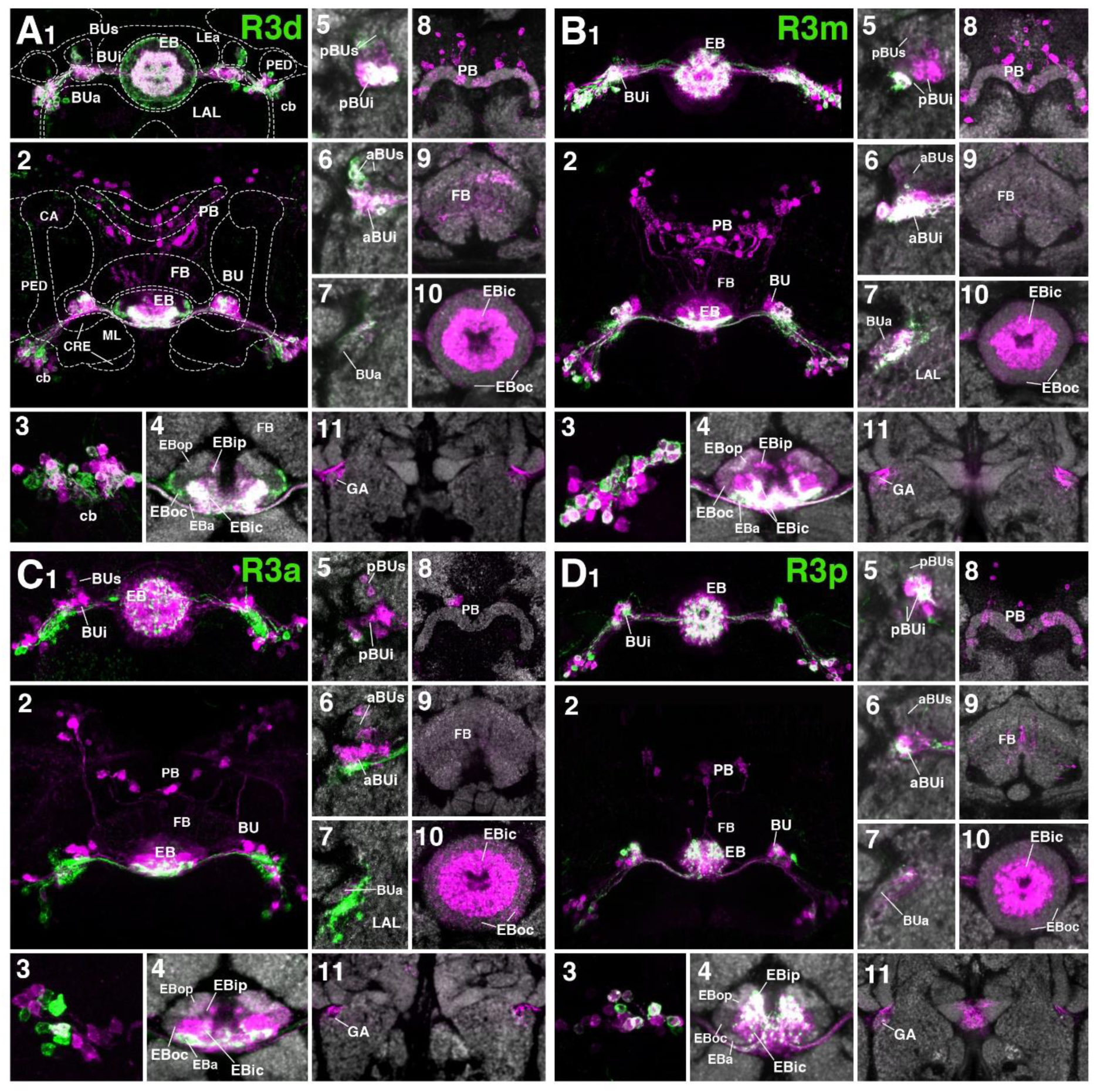
Putative postsynaptic partners of R-neurons revealed by *trans*-Tango (A-D) Similar modular format of panels as described in legend for Fig. 8. (A1-11) (R3d) R80C07-Gal4 > *trans*-Tango (B1-11) (R3m) R28E01-Gal4 > *trans*-Tango (C1-11) (R3a) R12G08-Gal4 > *trans*-Tango (D1-11) (R3p) VT063949-Gal4 > *trans*-Tango Other abbreviations: CRE, crepine; ML, PED, and CA, medial lobe, peduncle, and calyx of the mushroom body, respectively; BUa, BUs, and BUi, anterior, superior, and inferior domains of the bulb, respectively; anterior (EBa), inner central (EBic), outer central (EBoc), inner posterior (EBip), and outer posterior (EBop) domains of the ellipsoid body; LEa, anterior component of the lateral ellipsoid fascicle; cb, cell bodies.

### R3m

The pattern of trans-synaptically labeled neurons by the R3m driver 28E01-Gal4 largely resembles that described previously for R3d. R3m neurons have strong reciprocal interactions, with the majority of their cell bodies being positive for both GFP and RFP (Fig.9B1-B3). Based on the trans-synaptic labeling observed in EBic and EBip, as well as the periphery of EBoc (Fig.9B4), R3d, R3p and R4d neurons are also targeted by R3m. This is corroborated by labeling in the BU (Fig.9B5-B6), which includes pBUi (R3d), dorsal part of pBUi (R3p), and lateral part of BUs (R4d). Trans-synaptic labeling of E-PG neurons is sparse (Fig.9B8, B11).

**Figure 10.**
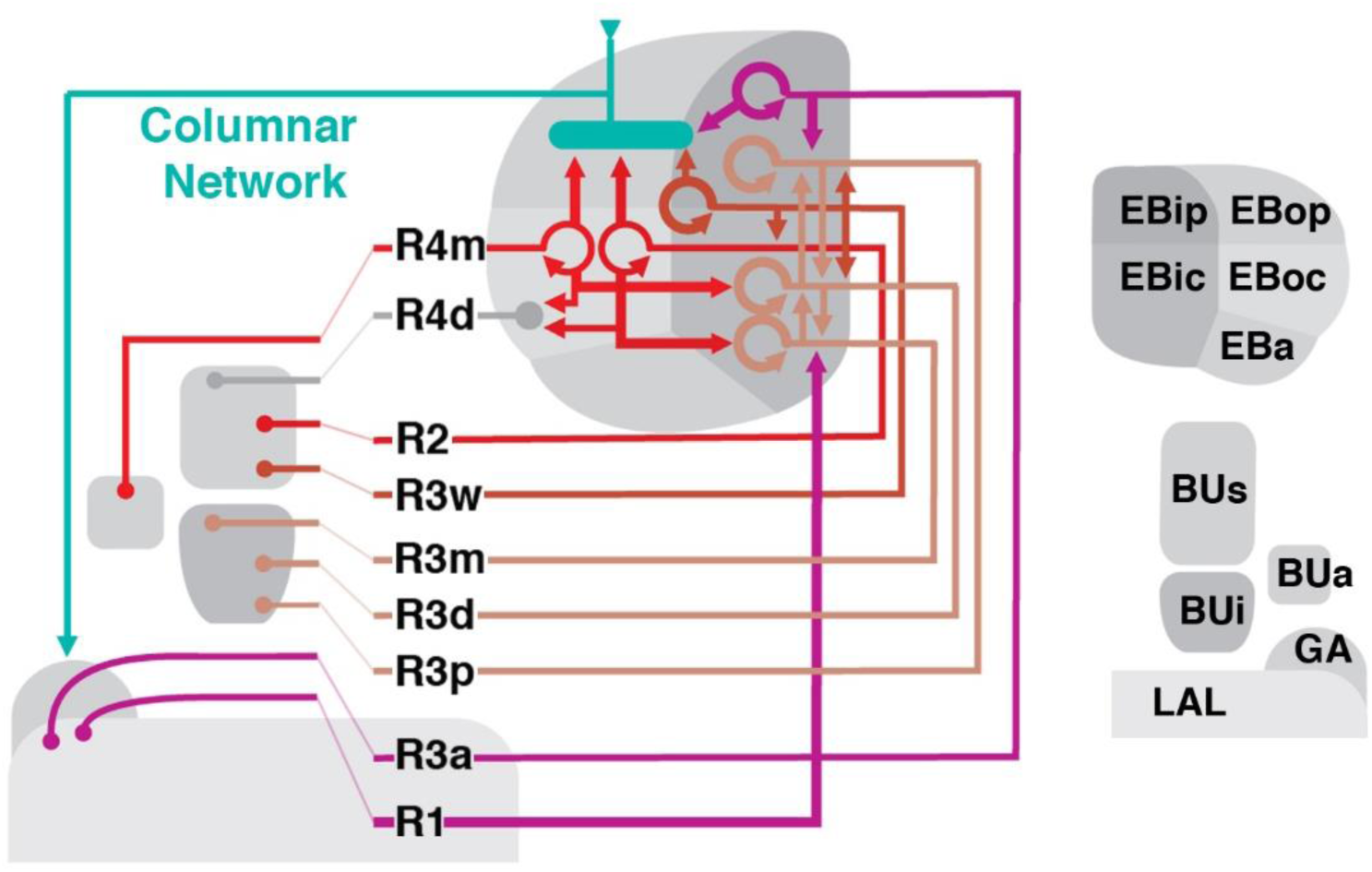
Schematized overview of putative R-neuron interactions.

### R3a

Whereas the inner R-neuron subclasses R3d, R3m and R3p preferentially form homotypic connections among each other, the R3a subclass targets EBic in a heterotypic manner. Thus, there is but little overlap of GFP and RFP in R-neuronal somata (Fig.9C1-C3). Strong target label is seen in EBic and throughout BUi (Fig.9C5-C6), indicating inner R-neuron subclasses R3d and R3m as preferred targets of R3a. Sparse label in EBoc and BUs suggests that some R2 neurons are also among the R3a targets (Fig.9C4-C6, C10). Furthermore, a very small number of E-PG neurons, and a matching faint RFP signal in EBop and EBoc, as well as the GA, appear to be targeted by R3a (Fig.9C2, C4, C11).

### R3p

R3p resembles R3d and R3m in forming strong homotypic interactions, as seen from overlapping GFP and RFP signal in cell bodies (Fig.9D1, D3), EBip (Fig.9D1-D2, D4) and dorsal part of pBUi (Fig.9D5). In addition, based on extensive RFP signal throughout BUi (Fig.9D5-D6), R3p targets other subclasses of inner R-neurons, notably R3d and R3m. We observed only small numbers of RFP-positive E-PG neuronal cell bodies (Fig.9D2, D8), matching only faint label of PB, outer EB, and GA (Fig.9D4, D8, D10-D11).

## Discussion

This work serves to build upon previous anatomical studies by further clarifying the neuronal architecture of the *Drosophila* ellipsoid body. Five definitive DN-cadherin domains constituting the EB neuropil provide fiducial landmarks with which neuron classes can be placed into spatial context. Based on this framework, we report several novel ring neuron subclasses and propose potential interactions between ring, columnar, and neuromodulatory neurons in the EB. Lastly, we experimentally mapped putative postsynaptic partners of R-neurons using *trans*-Tango, revealing insight into how information may be distributed throughout the EB and the rest of the CX. In addition to the neuroanatomical description of different populations, the identification of driver lines enables genetic access to label or manipulate these populations. This provides an entry point for future studies to probe the functional properties of each class and test the interactions proposed herein. In the following, we summarize the primary findings, speculate on the functional significance of CX wiring principles, and place our study into a developmental-neuroanatomical context with previous works in *Drosophila* and homologous structures in other insects.

### Information flow in the ellipsoid body network: input, recurrence, output, and neuromodulation

The CX is viewed as a critical hub for goal-directed navigational behavior in insects. Streams of sensory information from different modalities must converge onto this center of sensorimotor integration to guide navigational decisions based on current trajectory, learned information, and motivational state (Heinze, 2017). Central to this notion was the identification of a stable compass representation that tracks the flies heading in the E-PG neuron population. The robustness of this neural correlate of angular orientation, manifested as a single calcium activity “bump” that moves around the EB, depends on both visual and proprioceptive cues (Seelig and Jayaraman, 2015). Heavily relying upon studies in other insect species as a basis for comparison (Jundi et al., 2014), recent progress has been made towards identifying the neural pathways that transmit sensory information to the *Drosophila* CX, with visual input being the most well characterized. The fly CX receives visual information via the anterior visual pathway (AVP), a circuit defined by three successive layers. Information is transmitted from the optic lobe medulla to the anterior optic tubercle, from the tubercle to the bulb, and from there to the ellipsoid body, via medullo-tubercular (MeTu), tuberculo-bulbar (TuBu), and DALv2 ring neurons (R-neurons), respectively (Omoto et al., 2017). Parallel ensembles of TuBu neurons terminate in a topographically-organized fashion onto the microglomerular dendrites of distinct R-neuron subclasses within the bulb (Omoto et al., 2017). Specific computations are implemented across successive layers in this pathway, such as the integration of recent visual history and self-motion, which may inform downstream behavior (Shiozaki and Kazama, 2017; Sun et al., 2017). Ring neurons transmit processed visual information concerning features and landmarks to the EB, likely as a stable allothetic reference to guide bump dynamics in E-PG neurons (proposed in Seelig and Jayaraman, 2015; Turner-Evans et al., 2017). Indeed, we provide evidence that R2 neurons, which are tuned to visual features (Seelig and Jayaraman, 2013), provides direct presynaptic input to E-PG neurons (Fig.8B; Fig.10). The calcium activity bump in E-PG neurons also shift in total darkness, demonstrating the existence of a proprioceptive input channel that can update the heading representation in the EB in the absence of visual input. We posit that transmission of idiothetic cues to the CX is mediated in part by R1 and/or ExR4 neurons, as their neurite distribution and polarity suggests feedback from the LAL, a proposed motor signaling center (Fig.2; Supp. Fig.2; Fig.6) (Namiki and Kanzaki, 2016).

Conceivably, the information received by different R-neuron subclasses is transmitted to their ring-shaped neurites, and is processed via connections within the same subclass (homotypic interactions) and/or between subclasses (heterotypic interactions), the extent of which depends on the R-neuron subclass in question (Fig.8; Fig.9; Fig.10). As such, the R-neuron system likely displays recurrent connectivity to enable persistent activity required for memory processes, as has been shown for mushroom body circuits that support courtship memory (Zhao et al., 2018). Indeed, R3d (or R3c*) neurons, which comprise a critical nucleus of visual working memory (Neuser et al., 2008; Rieche et al., 2018), display prominent homotypic interactions (Fig.9A; Fig.10). Future work to define the mechanisms underlying intra-subclass interactions and experiments to perturb them, are required to assess the functional significance of these homotypic interactions.

R-neurons, particularly subclasses of which occupy peripheral EB domains, provide input to several different columnar neuron populations. To our knowledge, this study provides novel insight into the nature of subclass-specific, input-output communication between the ring and columnar networks. An important avenue of future work will be to elucidate the tuning properties of each R-neuron subclass and determine the contribution of each input to compass representation. Presumably, R-neuron subclasses that provide prominent, direct input to E-PG neurons, such as R2 or R4m (Fig.10), would exhibit the most influence over compass representation.

Circuit flexibility is likely facilitated by neuromodulatory input on a moment-by-moment basis, which may reconfigure information flow through the network and thus the output of the system. Neuromodulation would likely occur at multiple processing stages, as evidenced by the wide-spread neurites of dopaminergic neurons. For example, a single PPM3 neuron, innervates the GA/LAL, BU, and EBoc/op. We envisage that neurite-specific signaling and plasticity may regulate distinct processing nodes, akin to what has been demonstrated for dopaminergic neurons that encode protein hunger (Liu et al., 2017). Similarly, 5-HT may also influence R-neuron activity; projections from serotonergic neurons, likely from the posterior medial protocerebrum, dorsal cluster (PMPD; Pooryasin and Fiala, 2015), most prominently innervate EBic (Fig.5M). The effect of serotonin may be receptor and circuit specific; distinct 5-HT receptor isoforms are differentially expressed in specific R-neuron subclasses (Gnerer et al., 2015).

### Three-dimensional architecture of the ellipsoid body

For clarity, the five EB domains defined by the global marker DN-cadherin should be reconciled with previously used anatomical terminology of the EB (Omoto et al., 2017; Fig.1). Frontal sections of the EB at different anteroposterior depths shows that DN-cadherin domains are distinct, annular entities. These domains correspond to “layers” in other insects, and have sometimes been also referred to as layers in *Drosophila* as well (Young and Armstrong, 2010b). Therefore, N-cadherin EB domains are synonymous with layers. Each domain is best represented using a “dorsal standard view”: a horizontal section through the EB containing a lengthwise perspective of the EB canal (Fig.1). From this standard view, the N-cadherin domains are also clearly organized along the anteroposterior axis. Three anteroposterior subdivisions of the EB have been referred to as “shells”, in line with terminology used for the FB (Wolff et al., 2015). We propose that the anterior most shell encapsulates the anterior domain of the EB (EBa), and therefore consists of only one layer. The intermediate shell (called medial shell in Wolff et al., 2015) encapsulates the inner central (EBic) and outer central (EBoc) domains, and consists of two layers. Finally, the posterior shell encapsulates the inner posterior (EBip) and outer posterior (EBop) domains, and consists of two layers. For example, P-EN neurons occupy the EBop domain, which resides in the posterior EB shell (Fig.7D-E).

Lin et al., (2013) defined four substructures denoted as “rings” [EB_A_ (Anterior), EB_O_ (Outer), EB_C_ (Center), EB_P_ (Posterior)], which were based on anti-discs large (DLG) immunostaining and roughly correspond to the DN-cadherin domains. Like the DN-cadherin domains, each “ring” was proposed to contain specific R-neuron subclasses. Based on the ring neuron subclasses proposed by Lin et al., (2013) to comprise each “ring” (see below), we infer that EB_A_ from Lin et al. (2013) corresponds to EBa and EBic in our classification system. Furthermore, EB_O_ is EBoc, EB_C_ is EBip, and EB_P_ is EBop.

How does the annular domain structure of the *Drosophila* ellipsoid body compare to the lower division of the central body (CBL) described for other insects? Similar to the EB, the CBL represents a multilayered neuropil compartment formed by the neurite contributions of tangential and columnar elements. In insects such as locust (*Schistocerca gregaria*), which will be used as the primary basis for comparison in the following, the kidney bean or sausage-shaped CBL corresponds to the torus-shaped ellipsoid body in *Drosophila* (Ito et al., 2014; Pfeiffer and Homberg, 2014). In locusts, the CBL is effectively located ventrally of the upper division of the central body (CBU), whereas the homologous structures in *Drosophila* (ellipsoid body and fan-shaped body, respectively) are arranged in an antero-posterior fashion. This difference is reflective of a 60° anterior tilt of the locust neuraxis, as evidenced by the peduncle, which extends horizontally in flies but is oriented almost vertically in the locust (Hadeln et al., 2018). In the dung beetle (*Scarabaeus lamarcki*) and monarch butterfly (*Danaus plexippus*), the CBL are also sausage-shaped, but the neuraxis orientation is like that of *Drosophila* (Heinze and Reppert, 2012; Immonen et al., 2017). Differences in neuraxis orientation influence the comparison between the internal architecture of the locust CBL and fly EB. The locust CBL is subdivided along the dorso-ventral axis into 6 horizontal layers (although not stacked seamlessly on top of one another). Based on the expression of global markers, the *Drosophila* ellipsoid body is divided into toroidal domains (EBa/ic/oc/ip/op; Fig.1). Considering the tilt in neuraxis, we posit that dorsal strata (layers 1-2) of the locust CBL roughly correspond to more posterior domains (EBip/op) of the fly EB, whereas ventral strata (layers 3-6) correspond to more anterior EB domains (EBa/ic/oc). Corroborating this notion is the fact that fly P-EN neurons innervate EBop, and the locust homologues (called CL2 neurons) innervate dorsal layers of the CBL (Müller et al., 1997).

### Lineage-based architecture of the ellipsoid body

The EB and its domains, as well as other structures of the CX, are established by the neurite contributions of distinct neuronal populations. How is the neuronal diversity and connectivity of the CX developmentally established? The CX, and brain in general, is organized into structural-genetic modules called lineages; a lineage comprises the set of sibling neurons derived from an individual neural progenitor (neuroblasts). Each neuroblast forms a spatially discrete cluster of neurons with shared wiring properties; sibling neurons extend a limited number of fasciculated axon tract(s) and innervate specific brain compartments. Most brain lineages are “type I” neuroblast lineages, whose neuroblasts undergo a series of asymmetric divisions each of which renews the neuroblast and produces a ganglion mother cell. Columnar neurons of the CX are generated from four type II lineages which are larger and more complex than type I, with neuroblasts first producing a set of intermediate progenitors which in turn, give rise to ganglion mother cells (Bello et al., 2008; Boone and Doe, 2008; Bowman et al., 2008; Doe, 2017; Wang et al., 2014).

While the columnar neurons contributing to the EB are derived from type II lineages, the tangential elements (R-neurons) are largely derived from a single paired type I neuroblast, forming the lineage DALv2 (also called EBa1) (Ito et al., 2013; Omoto et al., 2017; Wong et al., 2013; Yu et al., 2013). Neurons of the DALv2 lineage have been studied in developmental contexts in a number of previous works (Kumar et al., 2009; Larsen et al., 2009; Lovick et al., 2017; Minocha et al., 2017; Spindler and Hartenstein, 2011; Xie et al., 2017). Production of secondary neurons by DALv2 begin around 24h after hatching (Lovick and Hartenstein, 2015). According to Kumar et al. (2009), one of the DALv2 hemilineages undergoes apoptotic cell death, implying that the DALv2 R-neurons forming the adult ellipsoid body represent a single hemilineage. Cursory heat-shock inducible single-cell clonal analysis carried out in the present study suggests that distinct R-neuron subclasses are born during specific time windows and therefore represent sublineages of DALv2 (Fig.4). Thus, clonal induction shortly after the onset of secondary neuroblast proliferation (20-48h after hatching) yielded exclusively outer R-neurons of the R4m subclass. At increasingly later time points, these types of clones become rare, and disappeared entirely at induction times after 96h. The converse is the case for inner ring neurons (R3d/m), which could be induced in increasing numbers with later time points of induction. Given that only a fraction of the overall number of R-neuron subclasses was represented among our clones, additional studies are required to settle the exact birth order of different R-neuron subclasses.

### Ontology of the ring neuron classification system

In the following, we provide a brief historical account of ring neuron definitions, attempt to resolve discrepancies in the literature when possible, and provide rationale for naming conventions used in this work.

Based on the description by Hanesch et al., (1989), the R-neuron type corresponds to ring neurons of the DALv2 lineage, with four R-neuron subclasses described in this initial study (R1-4). Two other ring neuron types were designated as “extrinsic ring neurons” (ExR-neurons), based on large projections outside of the EB; in this study, we pool neurons with this feature into a single type, the ExR-neurons. The first type of extrinsic R-neuron (the ExR1 subclass) described in Hanesch et al., (1989) likely corresponds to helicon cells. The second type (the ExR2 subclass) was not fully reconstructed by Hanesch et al., (1989), but due to its innervation of the caudal EB, ExR2 may correspond to the EBop-innervating PPM3 dopaminergic neuron (Fig.5N-Q), and thus our rationale for this designation. The serotonergic neurons that innervate the EB, likely the PMPD neurons, we designate as ExR3. Therefore, ExR1-3 are posteriorly localized ExR-neurons, likely deriving from the DM4-6 lineages. Due to its wide arborization and non-DALv2 based origin, we designate ring neurons of lineage BAmv1, with perikarya in the anterior cortex, as a fourth type of ExR-neuron (ExR4); we cannot exclude the possibility that ExR2 from Hanesch et al., (1989) may correspond to ExR4-neurons, as they too innervate the caudal EB. Furthermore, the “P”-neurons, described in Lin et al. (2013) as having ventrally localized cell bodies and also innervate the caudal EB, likely correspond to what we designate as ExR4-neurons.

Renn et al., (1999) was the first to conduct a genetic analysis of the EB neuropil, using enhancer-trap technology. Renn et al. (1999) described Line c105 to label R1 neurons (Supp. Fig.1A), due to their “inside-out” arborization pattern, inner ring localization, and extension into the posterior layers of the EB, a similar description to that of Hanesch et al. (1989). However, c105-positive R1 neurons exhibit ventrally projecting neurites into the LAL and lack bulb microglomeruli (Renn et al., 1999; Supp. Fig.1A3-5), in contrast to what was defined as R1 in Hanesch et al., (1989). Therefore, we speculate that Renn et al., (1999) identified a novel R-neuron class, distinct from R1 as described by Hanesch et al., (1989). However, due to R1 being the predominant designation this R-neuron subclass thereafter (Lin et al., 2013; Renn et al., 1999; Young and Armstrong, 2010b), we retain this classification in our study.

If R1 from Renn et al., (1999) was a previously undescribed class, what was R1 from Hanesch et al., (1989)? Based on the camera lucida drawn Golgi stained preparations and the description of R1 being “restricted to the inner zone lining the EB canal”, we propose that R1 from Hanesch et al., (1989) was interpreted by Renn et al., (1999) as R3. Henceforth, the predominant description of R3, which has been heavily investigated for their role in visual working memory, is that of “inner ring” R-neurons, a convention we also therefore retain (Kuntz et al., 2012; Lin et al., 2013; Neuser et al., 2008; Omoto et al., 2017; Young and Armstrong, 2010b). However, it is unclear which of the several R3-neuron subclasses with bulb microglomeruli identified in this study (R3m, R3d, R3p, or R3c*), correspond to R1 described in Hanesch et al. (1989).

Presuming R1 being renamed R3 by later authors, which R-neuron subclass corresponds to R3 in Hanesch et al., (1989)? R3 was shown to have an inside-out innervation pattern, restricted to the rostral half of the EB, and importantly, exhibited branches with terminals in the inner and outer ring (Hanesch et al., 1989). To our knowledge, this R-neuron subclass has not been described in any study thereafter, and likely corresponds to R3w of the present study (Fig.2G).

R4 was described by Hanesch et al., (1989) to project in an “outside-in” fashion and extend terminals into the outermost zone. Renn et al. (1999), noted two neuron subclasses that exhibit these features, and referred to these “R4-type” neurons as R4m and R4d. The camera lucida drawing of R4 in Hanesch et al., (1989) displays a ventrally localized microglomerulus (likely corresponding to BUa), a description that matches that of R4m in subsequent studies (Renn et al., 1999); we postulate that R4d was a class that exhibited R4-like wiring properties, newly identified by Renn et al., (1999) altogether.

R2 in former studies [Hanesch et al., 1989; labeled by c42 (Supp. Fig.1B; Renn et al., 1999) and EB1-Gal4 (Supp. Fig.1C; Young and Armstrong, 2010b) has axons within the EBoc domain, along with R4m (Fig.3B5-B6). In a more recent paper (Lin et al., 2013), the designation “R2” became associated with neurons projecting to the “anterior ring” (synonymous with EBa from this study). The designation of these anterior EB R-neurons as “R2” has carried forward to other studies, when their critical role in the regulation of sleep homeostasis was identified (Donlea et al., 2018; Liu et al., 2016). It should be noted that Liu et al., (2016) documented the distinction between what had historically been referred to as R2, and what had been referred as “R2” in Lin et al., (2013). “R2” in Lin et al. (2013) appears morphologically similar to a subclass revealed by the 52y driver in Young and Armstrong (2010), which were not given a specific name. We used the name R5 for these anterior R-neurons (Omoto et al., 2017) and propose to retain this designation to prevent future studies from equating them with what has been historically referred to as R2.

In more recent studies, the driver 38H02-Gal4 has been described as labeling R4 (or an R4-subset), in several studies (Dus et al., 2013; Ofstad et al., 2011; Park et al., 2016). 38H02-Gal4 does in fact label R4m (based on BUa microglomeruli and “outside-in” EBoc innervation pattern), but also strongly labels R5 (Supp. Fig.1J). Two other drivers, 15B07-Gal4 and 28D01-Gal4, were used to target EB neurons required for visual-thermal associations in place learning (Ofstad et al., 2011), and were described as labeling “R1 and R4”, or “R1 alone”, respectively. Anatomical re-assessment of these drivers reveals that 15B07-Gal4 labels R3d and R3p (or R3c*) and R4d (Supp. Fig.1I), whereas 28D01-Gal4 labels a neuron subclass indicative of R3m (Supp. Fig.1D).

In summary, the dorsal view of the EB in conjunction with DN-cadherin immunostaining provide criteria to more definitively identify ring neuron subclasses for future studies. The model organism *Drosophila* offers unique advantages to examine the circuit motifs that support the broadly relevant computations underlying the processes attributed to the CX; 1) the neurons comprising the CX are spatially and numerically confined, 2) genetic access to label, assess connectivity between, or functionally manipulate, specific neuron types within it, and 3) amenability to electro- or optophysiological recordings, oftentimes in the behaving animal. To fully leverage these advantages, we provide a systematic description of the ring neuron subclasses comprising the EB, genetic tools to access them, and provide insight into their interactions with other neurons of the CX.

## Author contribution statement

Conceptualization, JO; Methodology, JO, BN; Investigation, JO, BN, PK, JL; Writing – Original Draft, JO, VH; Writing – Review and Editing, JO, BN, PK, JD, VH; Visualization, JO, BN, PK, VH; Supervision and Funding Acquisition – JD, VH

## Acknowledgments

We thank the Bloomington Stock Center and the Developmental Studies Hybridoma Bank for fly strains and antibodies. We thank the following labs for kindly providing fly lines: lab of Gilad Barnea for *trans*-Tango flies (Talay et al., 2017), Mark Frye for TPH-Gal4 (Park et al., 2006), Heinrich Reichert for Poxn-Gal4 (Boll and Noll, 2002), and Barry Dickson for Vienna Tiles driver lines (Tirian and Dickson, 2017). Images from FlyCircuit were obtained from the NCHC (National Center for High-performance Computing) and NTHU (National Tsing Hua University), Hsinchu, Taiwan. This work was supported by the NIH (grants R01 NS096290 to V.H. and R01NS105967 to J.M.D.), additional support provided by the University of California, Los Angeles Dissertation Year Fellowship and the A.P. Giannini Postdoctoral Fellowship (to J.J.O.).

